# Dengue antigenic relationships predict evolutionary dynamics

**DOI:** 10.1101/432054

**Authors:** Sidney Bell, Leah Katzelnick, Trevor Bedford

## Abstract

Dengue virus (DENV) exists as four genetically distinct serotypes, each of which is historically assumed to be antigenically uniform. However, recent analyses suggest that antigenic heterogeneity may exist within each serotype, but its source, extent and impact remain unclear. Here, we construct a sequence-based model to directly map antigenic change to underlying genetic divergence. We identify 49 specific substitutions and four colinear substitution clusters that contribute to dengue antigenic diversity. We report moderate antigenic diversity within each serotype, resulting in variation in genotype-specific patterns of heterotypic cross-neutralization. We also quantify the impact of this antigenic heterogeneity on real-world DENV population dynamics. We find that antigenic fitness mediates fluctuations in DENV clade frequencies, although this appears to be primarily explained by coarser serotype-level antigenic differences. These results provide a more nuanced understanding of dengue antigenic evolution, with important ramifications for vaccine design and epidemic preparedness.

**Author Summary:** Dengue virus (DENV), the causative agent of dengue hemorrhagic fever, exists as four genetically distinct serotypes, DENV1 to DENV4. These serotypes are antigenically distinct: symptomatic reinfection with a homotypic virus is very rare, while reinfection with a heterotypic virus is sometimes associated with severe disease. Until recently, it has been assumed that viruses within each serotype are antigenically uniform. However, specific genotypes within each serotype have been anecdotally associated with varying severity of patient outcomes and epidemic magnitude. One hypothesis is that each serotype contains overlooked, meaningful antigenic diversity. While antigenic cartography conducted on neutralization titers suggests that heterogeneity may exist within each serotype, its source, extent and impact is unclear. Here, we analyze a previously published titer dataset to quantify and characterize the extent of DENV intraserotype antigenic diversity. We map antigenic changes to specific mutations in *E*, the dengue envelope protein, and interpolate across the alignment to estimate the antigenic distance between pairs of viruses based on their genetic differences. We identify 49 specific substitutions and four colinear substitution clusters that contribute to dengue antigenic evolution. We find that DENV antigenic divergence is tightly coupled to DENV genetic divergence, and is likely a gradual, ongoing process. We report modest but significant antigenic diversity within each serotype of DENV, which may have important ramifications for vaccine design. To understand the impact of this antigenic heterogeneity on real-world DENV population dynamics, we also quantify the extent to which population immunity—accumulated through recent circulation of antigenically similar genotypes—determines the success and decline of DENV clades in a hyperendemic population. We find that antigenic fitness is a key determinant of DENV population turnover, although this appears to be driven by coarser serotype-level antigenic differences. By leveraging both molecular data and real-world population dynamics, these results provide a more nuanced understanding of dengue antigenic evolution, with important ramifications for improving vaccine design and epidemic preparedness.

## Introduction

Dengue virus (DENV) is a mosquito-borne flavivirus which consists of four genetically distinct clades, canonically thought of as serotypes (DENV1 – DENV4) (Lanciotti et al., 1997). DENV circulates primarily in South America and Southeast Asia, infecting 400 million people annually. Primary DENV infection is more often mild and is thought to generate lifelong homotypic immunity and temporary heterotypic immunity, which typically wanes over six months to two years (Katzelnick et al., 2016; Reich et al., 2013; Sabin, 1952). Subsequent heterotypic secondary infection induces broad cross-protection, and symptomatic tertiary and quaternary cases are rare (Gibbons et al., 2007; Olkowski et al., 2013). However, a small subset of secondary infections are enhanced by non-neutralizing, cross-reactive antibodies, resulting in severe disease via antibody dependent enhancement (ADE) (Halstead, 1979; Katzelnick et al., 2017; Salje et al., 2018; Sangkawibha et al., 1984). Approximately 1–3% of cases progress to severe dengue hemorrhagic fever, causing ~9,000 deaths each year (Bhatt et al., 2013; Stanaway et al., 2016) and relative risk of severe dengue from secondary heterotypic infection relative to primary infection is estimated to be ~24 (Mizumoto et al., 2014). Thus, the antigenic relationships between dengue viruses — describing whether the immune response generated after primary infection results in protection or enhancement of secondary infection — are key drivers of DENV case outcomes and epidemic patterns.

While each serotype is clearly genetically and antigenically distinct, it is not clear how subserotype clades of DENV interact antigenically. Each DENV serotype consists of broad genetic diversity (Figure 1A), including canonical clades termed ‘genotypes’ (Rico-Hesse, 1990; Twiddy et al., 2003). Specific genotypes have been associated with characteristically mild or severe disease, and heterogeneous neutralization titers suggest that the immune response to some genotypes is more cross-protective than others (Gentry et al., 1982; Russell and Nisalak, 1967). Until recently, it has been assumed that these intraserotype differences are minimally important compared to interserotype differences. However, empirical evidence has demonstrated that these genotype-specific differences can drive case outcomes and epidemic severity (reviewed in Holmes and Twiddy (2003)). For example, analysis of a longitudinal cohort study demonstrated that specific combinations of primary infection serotype and secondary infection genotype can mediate individual case outcomes (OhAinle et al., 2011). On a population scale, the DENV1-immune population of Iquitos, Peru, experienced either entirely asymptomatic or very severe epidemic seasons in response to two different genotypes of DENV2 (Kochel et al., 2002).

**Figure 1.**
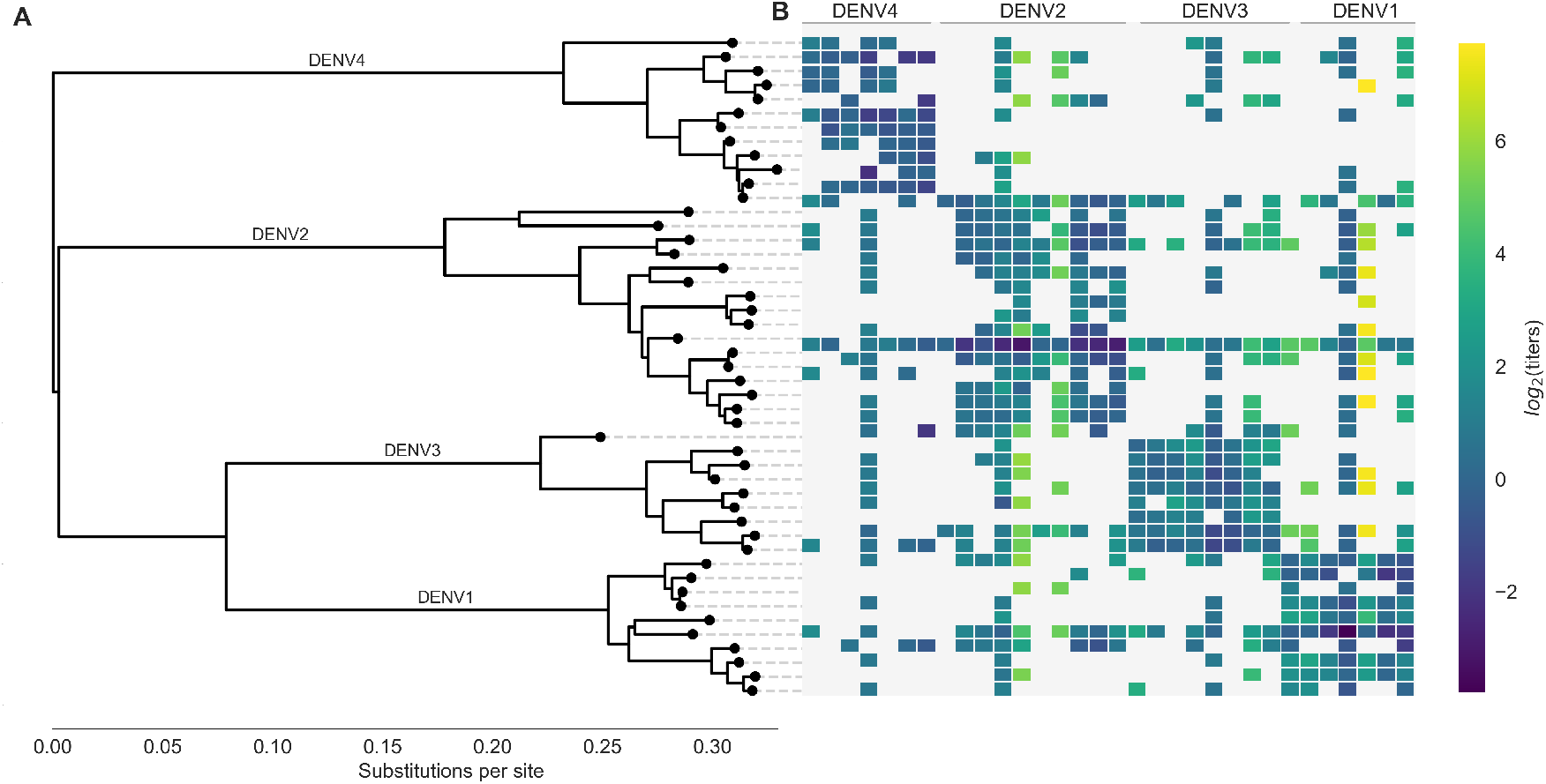
Phylogeny of dengue virus sequences and normalized antigenic distances. **(A)** Maximum likelihood phylogeny of the *E* (envelope) gene from titered dengue viruses. Notably, each of the four serotypes contains substantial genetic diversity. **(B)** Pairwise antigenic distances were estimated by Katzelnick *et al*. using plaque reduction neutralization titers (PRNT50, see Methods). Aggregated titer values are standardized such that the distance between autologous virus-serum pairs is 0, and each titer unit corresponds to a two-fold change in PRNT50 value. Light gray areas represent missing data. Larger values correspond to greater antigenic distance.

One explanation for these and similar observations is that overlooked intraserotype antigenic variation contributes to these genotype-specific case outcomes and epidemic patterns. Recent efforts to antigenically characterize diverse DENV viruses suggests that each serotype may contain antigenic heterogeneity, but the source and impact of this heterogeneity is not clear (Katzelnick et al., 2015). Here, we characterize the evolutionary basis for observed antigenic heterogeneity among DENV clades. We also quantify the impact of within- and between-serotype antigenic variation on real-world DENV population dynamics.

## Results

### Dengue neutralization titer data

Antigenic distance between a pair of viruses *i* and *j* is experimentally quantified using neutralization titers, which measure how well serum drawn after infection with virus *j* is able to neutralize virus *i in vitro* (Russell and Nisalak, 1967). Throughout the following we refer to serum raised against virus *j* as serum *j* for brevity. To measure the pairwise antigenic distances for a panel of diverse DENV viruses (Figure 1), Katzelnick *et al*. infected naive non-human primates (NHP) with each virus, drew sera at three months post-infection, and then titered this sera against a panel of test viruses (Katzelnick et al., 2015). To compare patterns of cross-protection in NHP and humans, they also drew sera from 31 study participants six weeks after inoculation with a monovalent component of the NIH dengue vaccine candidate. This sera was also titered against a broad panel of DENV viruses. As originally reported, we find generally consistent patterns of neutralization between the NHP and human sera data; see Katzelnick et al. (2015) for a detailed comparison. In total, our dataset consists of 454 NHP sera titers spanning the breadth of DENV diversity, and 728 human sera titers providing deep coverage of a small subset of viruses.

To normalize these measurements, we take the log_2_ of each value, such that one antigenic unit corresponds to a two-fold drop in neutralization, and we define antigenic distance between autologous serum-virus pairs (i.e., virus *i* and serum *i*) as zero. Normalized antigenic distance between virus *i* and serum *j* is thus calculated as *D_ij_* = log_2_(*T_ii_*) – log_2_(*T_ij_*), such that a higher value of *D_ij_* indicates that serum *j* is less effective at neutralizing virus *i*, implying greater antigenic distance between viruses *i* and *j*. For brevity, these normalized titer values are hereafter referred to simply as log_2_(titers).

The full dataset of standardized titer values is shown in Figure 1B. Here, we see that homotypic virus-serum pairs are more closely related antigenically than heterotypic pairs. However, we also observe large variance around this trend, both within and between serotypes. This suggests that treating each serotype as antigenically uniform potentially overlooks important antigenic heterogeneity across viruses within each serotype.

### Dengue antigenic evolution corresponds to genetic divergence

Titer measurements are prone to noise, and there is a limited amount of available titer data. If the antigenic heterogeneity observed in the raw data is truly the result of an underlying evolutionary process, we expect that differences in antigenic phenotype correspond to underlying mutations in surface proteins. Dengue has two surface proteins, prM (membrane) and E (envelope). While previous studies have identified epitopes on both prM and E, it is believed that antibodies involved in ADE primarily target prM, while neutralizing antibodies primarily target E (de Alwis et al., 2014). The assay used to generate this titer dataset captures neutralization, but does not capture the effects of ADE; we thus focus our analysis on the *E* gene.

To fully map the relationship between DENV genetic and antigenic evolution, we adapt a substitution-based model originally developed for influenza (Neher et al., 2016). Conceptually, this model predicts titer values through three steps. First, we align *E* gene sequences from titered dengue viruses and catalog the amino acid mutations between each serum strain and test virus strain in our dataset. Next, we infer how much antigenic change is attributable to specific mutations by constructing a parsimonious model that links normalized antigenic distances to observed mutations. This assigns each mutation *m* an antigenic effect size, *d_m_* ≥ 0; forward and reverse mutations are assigned separate values of *d_m_*. With this in hand, we estimate the asymmetrical antigenic distance 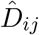 between all pairs of sera and test viruses by summing over *d_m_* for all mutations observed between the serum and the test virus (Methods, Eq. 2).

To learn these values of *d_m_*, we first split our dataset into training (random 90% of measurements) and test data (the remaining 10% of values). We take the training data and fit *d_m_* for each mutation that is observed two or more times, subject to regularization as follows (also detailed in Methods, Eq. 3). Parsimoniously, we expect that antigenic change is more likely to be incurred by a few key mutations than by many mutations; correspondingly, our prior expectation of values of *d_m_* is exponentially distributed such that most values of *d_m_* = 0. This is directly analogous to lasso regression to identify a few parameters with positive weights and set other parameters to 0 (Tibshirani, 1996). Additionally, some viruses have greater binding avidity, and some sera are more potent than others (Figure S1); these ‘row’ and ‘column’ effects, respectively, are normally distributed and are taken into account when training the model. The model uses convex optimization to learn the values of *d_m_* that minimize the sum of squared errors (SSE) between observed and predicted titers in the training data. We thus learn model parameters from the training data, and then use those parameters to predict test data values. We assess model performance by comparing the predicted test titer values to the actual values, aggregated across 100-fold Monte Carlo cross validation.

This model formulation is an effective tool for estimating antigenic relationships between viruses based on their genetic sequences. On average across cross-validation replicates, this model yields a root mean squared error (RMSE) of 0.75 when predicting titers relative to their true value (95% CI 0.74–0.77, RMSE), and explains 78% of the observed variation in neutralization titers overall (95% CI 0.77–0.79, Pearson *R*^2^). This is comparable to the model error from a cartography-based characterization of the same dataset (RMSE 0.65–0.8 log_2_ titer units) (Katzelnick et al., 2015). Prediction error was comparable between human and non-human primate sera, indicating that these genetic determinants of antigenic phenotypes are not host species-specific (Figure S2).

The 48 strains included in the titer dataset (as serum strains, test virus strains, or both) are 25.7% divergent on average (amino acid differences in *E*). Pairwise comparisons of all serum strains and test viruses yields 1,534 unique mutations that are observed at least twice. Our parsimonious model attributes antigenic change to a total of 49 specific mutations and 4 colinear mutation clusters (each consisting of 2–6 co-occurring mutations) (Figure 2, Table S1). Each of these mutations confers 0.01–2.11 (median 0.19) log_2_ titer units of antigenic change; 27 mutations or mutation clusters have *d_m_* ≥ 0.2. These mutations span all domains of *E*, and most occur both between and within serotypes (Figure 2).

**Figure 2.**
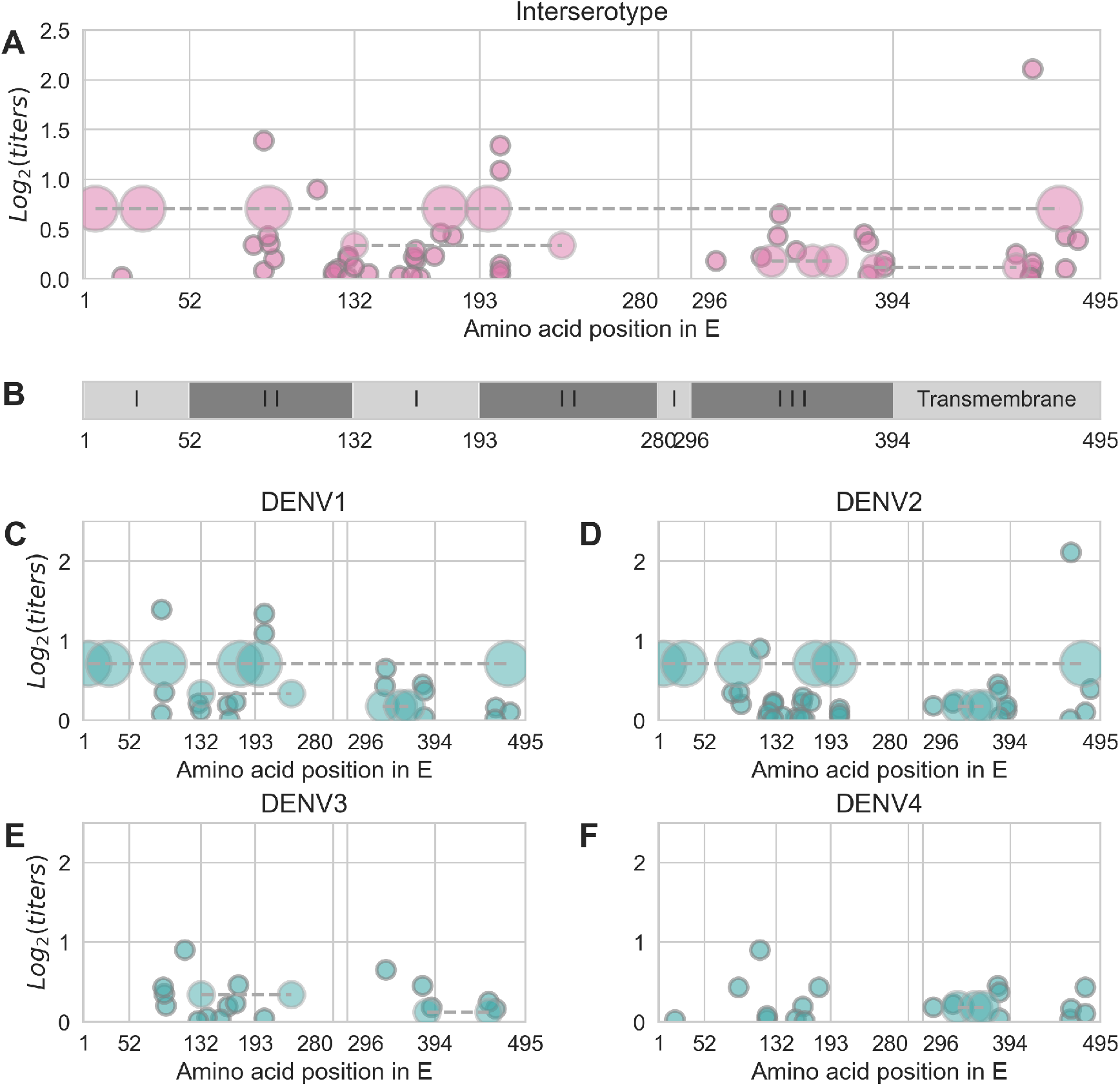
Distribution and effect size of antigenic mutations. Each point represents one antigenically relevant mutation or colinear mutation cluster. Clustered mutations are connected with dashed lines with point size proportionate to cluster size (N=2–6). The x axis indicates mutations’ position in *E*, relative to each functional domain as noted in **(B)**. The y axis indicates antigenic effect size.

### Each serotype of dengue contains moderate antigenic heterogeneity

By linking antigenic change to specific mutations, we are able to estimate unmeasured antigenic distances between any pair of viruses in the dataset based on their genetic differences. As an example, we estimated the antigenic distance between serum raised against each monovalent component of the NIH vaccine candidate and all other viruses in the dataset. As shown in Figure 3, vaccine-elicited antibodies result in strong homotypic neutralization, but heterotypic cross-neutralization varies widely between specific strains. This has important ramifications for vaccine design and trial evaluation.

**Figure 3.**
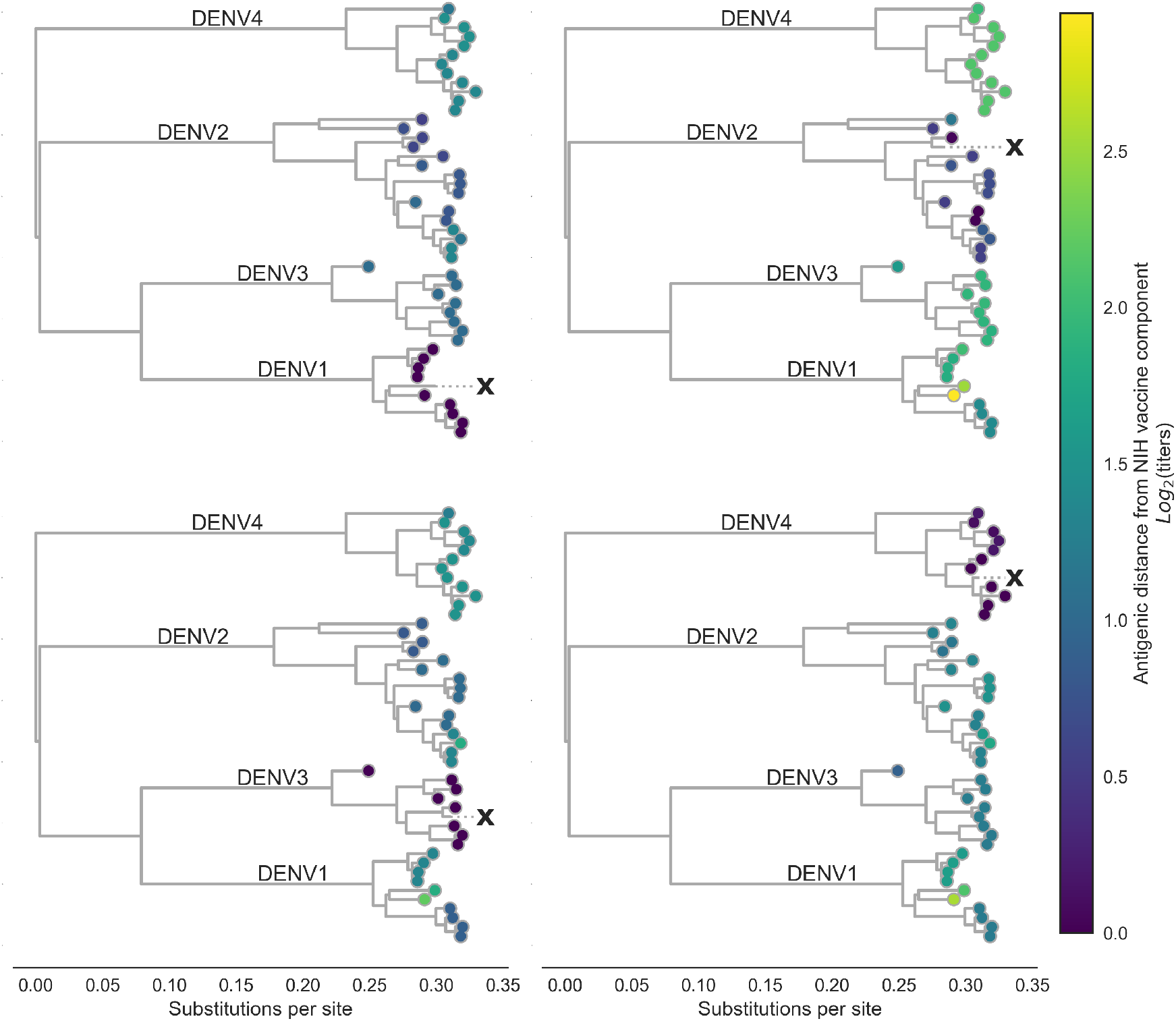
Antigenic distance from NIH vaccine strains. By assigning a discrete increment of antigenic change to each mutation, we can estimate the asymmetrical antigenic distance between any serum strain and test virus strain based on their genetic differences. Here, we show the estimated antigenic distance between serum raised against each monovalent component of the NIH vaccine candidate (indicated as ‘X’) and each test virus in the tree.

We also observe antigenic heterogeneity at the genotype level. On average, heterotypic genotypes are separated by 6.9 antigenic mutations (or colinear mutation clusters) and 2.18 log_2_ titers. Homotypic genotypes are separated by a mean of 1.9 antigenic mutations, conferring a total of 0.30 log_2_ titers of antigenic distance (Figure 4). Notably, the titer dataset spans the breadth of canonical DENV genotypes, but in most cases lacks the resolution to detect within-genotype antigenic diversity. We thus expect that these results represent a lower-bound on the true extent of DENV intraserotype antigenic diversity.

**Figure 4.**
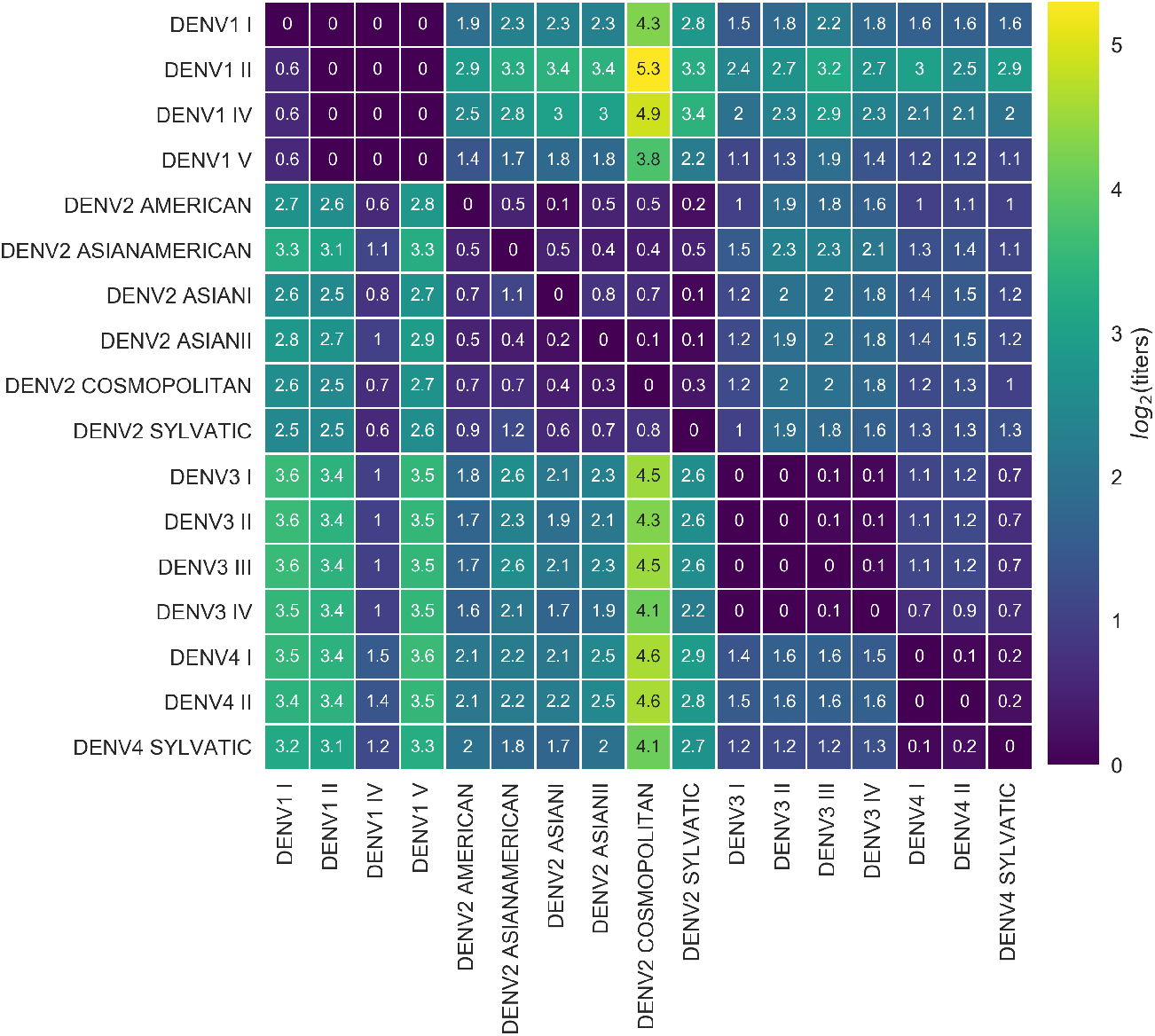
Titer distance by genotype. Values represent the mean interpolated antigenic distance between canonical dengue genotypes (in standardized log_2_ titer units).

In summary, we have identified a small number of antigenically relevant mutations that explain most of the observed antigenic heterogeneity in dengue, as indicated by neutralization titers. These mutations occur both between and within serotypes, suggesting that dengue antigenic evolution is an ongoing, though gradual, process. This results in strain-specific and genotype-level antigenic variation, although the scale of this variation is small compared to serotype-level differences. From this, we conclude that there is antigenic variation within each serotype of DENV, and that this is driven by underlying genetic divergence.

### Antigenic novelty predicts serotype success

From the titer model, we find evidence that homotypic genotypes of DENV vary in their ability to escape antibody neutralization. However, antibody neutralization is only one of many factors that shape epidemic patterns. We investigate whether the observed antigenic diversity influences dengue population dynamics in the real world.

The size of the viral population (i.e., prevalence, commonly analyzed using SIR models as reviewed in Lourenço et al. (2018)) is determined by many complex factors, and reliable values for population prevalence are largely unavailable. Contrastingly, the composition of the viral population (i.e., the relative frequency of each viral clade currently circulating) can be estimated over time by examining historical sequence data (Lee et al., 2018; Neher et al., 2016), and is primarily driven by viral fitness (Bedford et al., 2011).

In meaningfully antigenically diverse viral populations, antigenic novelty (relative to standing population immunity) contributes to viral fitness: as a given virus *i* circulates in a population, the proportion of the population that is susceptible to infection with *i*–and other viruses antigenically similar to *i*–decreases over time as more people acquire immunity (Bedford et al., 2012; Łuksza and Lässig, 2014). Antigenically novel viruses that are able to escape this population immunity are better able to infect hosts and sustain transmission chains, making them fitter than the previously circulating viruses (Bedford et al., 2012; Gupta et al., 1998; Lourenço and Recker, 2013; Wearing and Rohani, 2006; Zhang et al., 2005). Thus, if antigenic novelty constitutes a fitness advantage for DENV, then we would expect greater antigenic distance from recently circulating viruses to correlate with higher growth rates.

To test this hypothesis, we examine the composition of the dengue virus population in Southeast Asia from 1970 to 2015. We estimate the relative population frequency of each DENV serotype at three month intervals, *x_i_* (*t*) (Figure 5A), based on their observed relative abundance in the ‘slice’ of the phylogeny corresponding to each timepoint (N=8,644 viruses; see Methods, Eq. 4). While there is insufficient data to directly compare these estimated frequencies to regional case counts, we see good qualitative concordance between frequencies similarly estimated for Thailand and previously reported case counts from Bangkok (Figure S3).

**Figure 5.**
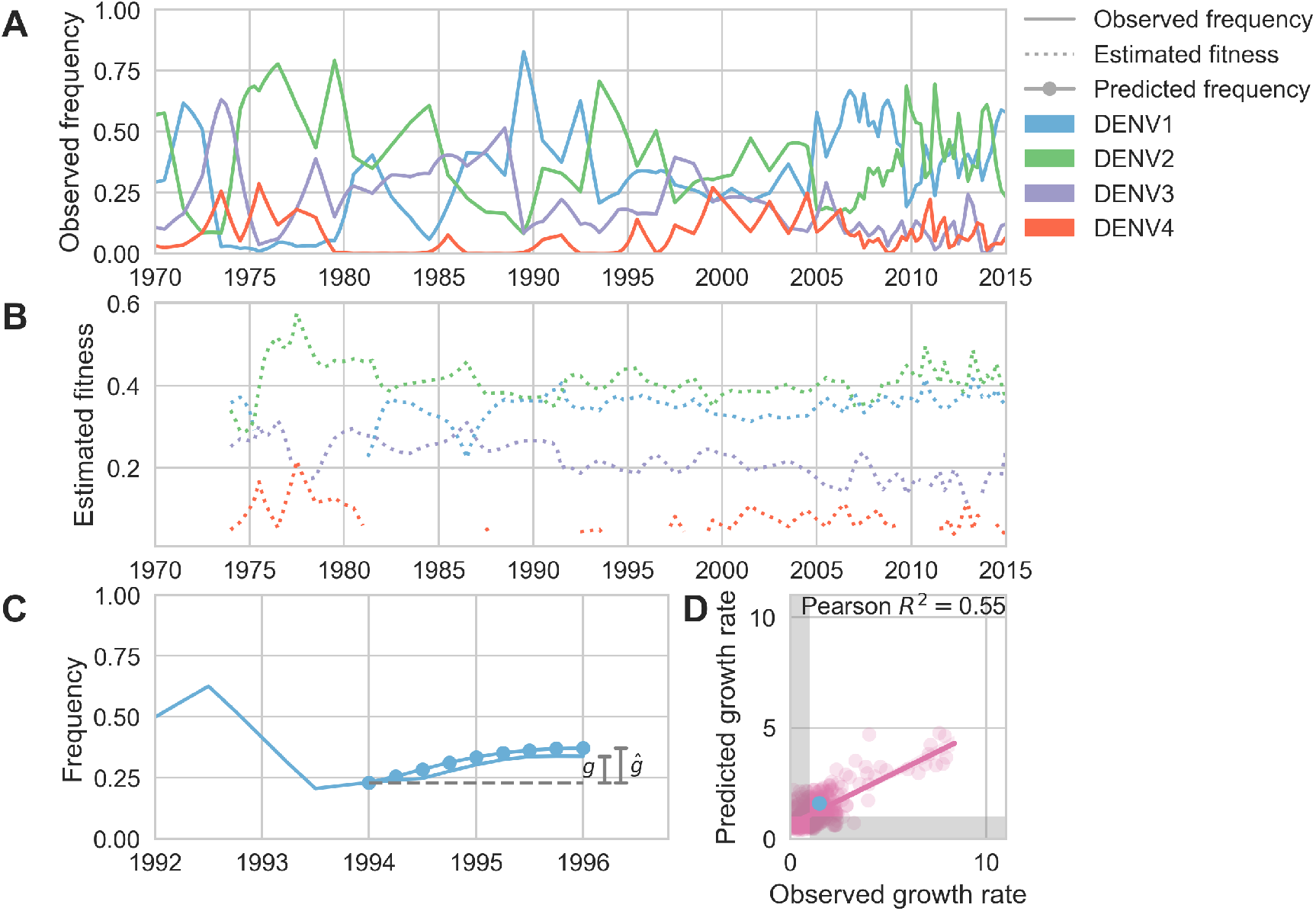
Antigenic novelty predicts serotype success. **(A)** The relative frequency of each serotype, *x_i_*, in Southeast Asia estimated every three months based on available sequence data. **(B)** Total fitness of each serotype. We calculate antigenic fitness for each serotype over time as its frequency-weighted antigenic distance from recently circulating viruses. We then add this to a time-invariant intrinsic fitness value to calculate total fitness. **(C)** DENV1 frequencies between 1994 and 1996 alongside model projection. At each timepoint *t*, we blind the model to all empirical data from timepoints later than *t* and predict each serotype’s future trajectory based on its initial frequency, time-invariant intrinsic fitness, and antigenic fitness at time *t* (Methods, Eq. 11). We predict forward in three-month increments for a total prediction period of *dt* = 2 years. At each increment, we use the current predicted frequency to adjust our estimates of antigenic fitness on a rolling basis (Methods, Eq. 15). **(D)** Predicted growth rates, 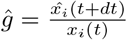, compared to empirically observed growth rates, 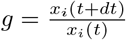. Predicted and empirical growth rate of the example illustrated in **(C)** is shown in **(D)** as the blue point. Serotype growth versus decline is accurate (i.e., the predicted and actual growth rates are both > 1 or both < 1, all points outside the gray area) for 66% of predictions.

Fitter virus clades increase in frequency over time, such that *x_i_* (*t* + *dt*) > *x_i_* (*t*). It follows that these clades have a growth rate—defined as the fold-change in frequency over time—greater than one: 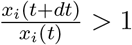. To isolate the extent to which antigenic fitness contributes to clade success and decline, we extend work by Łuksza and Lässig (2014) to build a simple model that attempts to predict clade growth rates based on two variables: the antigenic fitness of the clade at time *t*, and a time-invariant free parameter representing the intrinsic fitness of the serotype the clade belongs to. We estimate the antigenic fitness of clade *i* at time *t* as a function of its antigenic distance from each viral clade *j* that has circulated in the same population over the previous two years, weighted by the relative frequency of *j* and adjusted for waning population immunity (Figure 5B; Methods, Eq. 7). Growth rates are estimated based on a two year sliding window (Figure 5C).

This simple model explains 54.7% of the observed variation in serotype growth rates, and predicts serotype growth vs. decline correctly for 66.0% of predictions (Figure 5D). This suggests that antigenic fitness is a major driver of serotype population dynamics. This also demonstrates that this model captures key components of dengue population dynamics; examining the formulation of this model in more detail can yield insights into how antigenic relationships influence DENV population composition. The fitness model includes six free parameters that are optimized such that the model most accurately reproduces the observed fluctuations in DENV population composition (minimizing the RMSE of frequency predictions, see Methods). We find that serotype fluctuations are consistent with a model wherein population immunity wanes linearly over time, with the probability of protection dropping by about 63% per year for the first two years after primary infection. This model assumes no fundamental difference between homotypic and heterotypic reinfection; rather, homotypic immunity is assumed to wane at the same rate as heterotypic immunity, but starts from a higher baseline of protection based on closer antigenic distances. We also find that these dynamics are best explained by intrinsic fitness that moderately varies by serotype (Table 1); we are not aware of any literature that directly addresses this observation via competition experiments. However, intrinsic fitness alone is unable to predict serotype dynamics (Table S3) and relative strength of antigenic fitness and intrinsic fitness are approximately matched in determining overall serotype fitness.

**Table 1.** Optimized fitness model parameters for primary analysis

| Parameter | Value | Description |
| --- | --- | --- |
| $\beta$ | 1.02 | Slope of linear relationship between population immunity and viral fitness |
| $\gamma$ | 0.83 | Proportion of titers waning each year since primary infection |
| $\sigma$ | 0.76 | Slope of linear relationship between titers and probability of protection |
| $f_0^{(1)}$ | 0.74 | Relative intrinsic fitness of DENV1 |
| $f_0^{(2)}$ | 0.84 | Relative intrinsic fitness of DENV2 |
| $f_0^{(3)}$ | 0.50 | Relative intrinsic fitness of DENV3 |
| $f_0^{(4)}$ | 0.00 | Relative intrinsic fitness of DENV4 (fixed) |

### Antigenic novelty also partially predicts genotype success

To estimate how well antigenic fitness predicts genotype dynamics, we used the same model to predict genotype success and decline. As before, fitness of genotype *i* is based on the intrinsic fitness of the serotype *i* belongs to, and the antigenic distance between *i* and each other genotype, *j*, that has recently circulated (Figure 6B). For genotypes, we can calculate antigenic distance between *i* and *j* at either the serotype level or the genotype level. In the ‘interserotype model’, we treat each serotype as antigenically uniform, and assign the mean serotype-level antigenic distances to all pairs of constituent genotypes. In the ‘intergenotype model’, we incorporate the observed within-serotype heterogeneity, and use the mean genotype-level antigenic distances (as shown in Figure 4). If within-serotype antigenic heterogeneity contributes to genotype fitness, then we would expect estimates of antigenic fitness based on the ‘intergenotype model’ to better predict genotype growth rates.

**Figure 6.**
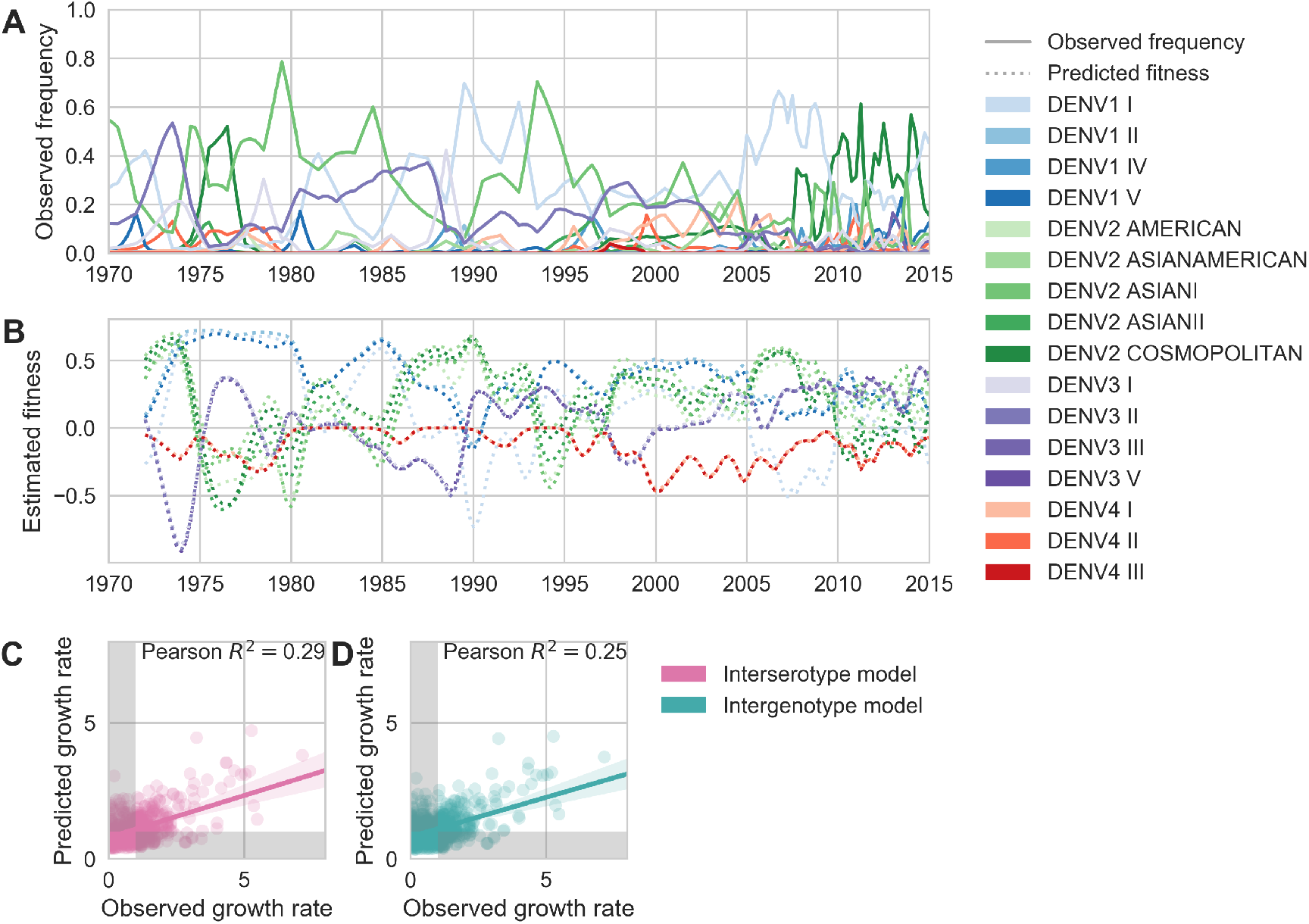
Antigenic novelty partially predicts genotype success. **(A)** Relative frequencies of each canonical dengue genotype across Southeast Asia, estimated from available sequence data. **(B)** Antigenic fitness is calculated for each genotype as its frequency-weighted antigenic distance from recently circulating genotypes. We then add this to a time-invariant, serotype-specific intrinsic fitness value to calculate total fitness (shown here, arbitrary units). We assess antigenic distance at either the ‘intergenotype’ or the ‘interserotype’ resolution. In this panel, we show total fitness over time, incorporating estimates of antigenic fitness derived from the ‘intergenotype’ model. **(C, D)** Fitness estimates were used to predict clade growth rates over 2 years, compounding immunity every three months based on predicted frequency changes (Methods Eq. 15). Here, we compare observed vs. predicted growth rates for both formulations of the fitness model (using fitness derived from either ‘interserotype’ or ‘intergenotype’ antigenic distances). Growth versus decline was accurate (predicted and actual growth rates both > 1 or both < 1, points outside the gray shaded area) for 67% and 61% of predictions, respectively.

We find that antigenic fitness contributes to genotype turnover, although it explains less of the observed variation than for serotypes. As for serotypes, intrinsic fitness alone was unable to predict genotype turnover (Table S3). When antigenic distance is estimated from the ‘interserotype model’, we find that our model of antigenic fitness explains approximately 28.6% of the observed variation in genotype growth rates, and correctly predicts genotype growth vs. decline 66.6% of the time (Figure 6C). Perhaps surprisingly, more precise estimates of antigenic distance between genotypes from the ‘intergenotype model’ does not improve our predictions of genotype success (*R*^2^ = 0.254, 61.0% accuracy; Figure 6D, Table S3). This suggests that although we find strong evidence that genotypes vary in their ability to escape neutralizing antibodies, these differences are subtle enough that they do not impact broad-scale regional dynamics over time.

## Discussion

### Within-serotype antigenic heterogeneity

We show that mapping antigenic change to specific mutations and interpolating across the DENV alignment is able to explain a large majority of the observed variation in antigenic phenotypes, as measured by neutralization titers. We identify 49 specific mutations and four colinear mutation clusters that contribute to antigenic variation, of which 27 mutations or mutation clusters have an antigenic impact of 0.20 log_2_ titers or greater. These mutations span all major domains of *E*, and occur both within and between serotypes. This demonstrates that DENV antigenic divergence is closely coupled to genetic divergence. We use these mutations to infer unmeasured antigenic relationships between viruses, revealing substantial within-serotype antigenic variation. For comparison, we reconstructed the ancestral sequence of each serotype and constrained the model to only permit antigenic change to be attributed to these serotype-level differences. While this interserotype-only model predicts titers to a reasonable degree, we find that it has higher error (RMSE = 0.86) than the full model which accounts for within-serotype heterogeneity (RMSE = 0.79; Table S2). This supports and expands upon previous reports (Forshey et al., 2016; Katzelnick et al., 2015; Waggoner et al., 2016) that the null hypothesis of antigenically uniform serotypes is inconsistent with observed patterns of cross-protection and susceptibility.

Consistent with the relatively long timescale of dengue evolution, we observe many sites in the dengue phylogeny to have mutated multiple times. These represent instances of parallelism, reversion and homoplasy. For example, we observe that site 390 is consistently S in DENV1, N in DENV3 and H in DENV4, while DENV2 genotypes show a mixture of D, N and S (S4). We estimate an antigenic impact of 0.18 log_2_ titers of the N390S mutation. Our model predicts that the parallel N390S mutations in DENV1 and DENV2 Cosmopolitan makes these viruses slightly more antigenically similar rather than more antigenically distinct. Along these lines, we compared the ‘substitution’ model to a similar model formulation (termed the ‘tree’ model) which assigns *d_m_* values to individual branches in the phylogeny, rather than to individual mutations, so that each branch with a positive *d_m_* value increases antigenic distance between strains (Neher et al., 2016). As expected from the high degree of homoplasy across the dengue phylogeny, we observe that the ‘substitution’ model outperforms the ‘tree’ model in predicting titers in validation datasets (Table S2).

To investigate the impact of this observed variation, we examine patterns of neutralization in response to vaccination with each monovalent component of the NIH vaccine candidate. Here, we see that each monovalent component elicits broad homotypic protection, but levels of heterotypic cross protection vary widely between heterotypic genotypes. This is consistent with previous reports of genotype-specific interactions between standing population immunity and subsequent heterotypic epidemics as modulating epidemic severity (Kochel et al., 2002; OhAinle et al., 2011). We hypothesize that this observed within-serotype variation primarily effects heterotypic secondary infection outcomes, rather than modulating homotypic immunity. Although we note that Juraska et al. (2018) demonstrate that vaccine efficacy decreases with increasing amino acid divergence of breakthrough infections from the vaccine insert.

Overall, we expect that these antigenic phenotypes represent a lower-bound on the extent, magnitude, and nature of antigenic heterogeneity with DENV. Our current titer dataset spans the breadth of DENV diversity, but due to small sample size, it lacks the resolution to detect most sub-genotype antigenic variation. The appearance of the deep antigenic divergence of the four serotypes, and the more recent antigenic divergences within each serotype, suggest that DENV antigenic evolution is likely an ongoing, though gradual, process. We therefore expect that future studies with richer datasets will find additional antigenic variation within each genotype. This dataset also contains many left-censored titer values, where we know two viruses are at least *T* titer units apart, but do not know exactly how far apart. If we knew the true value of these censored titers, many of them would indicate larger antigenic distances than the reported values, *T*, which are used to train the model. Thus, it is likely that our model systematically underestimates the magnitude of titer distances.

Finally, antibody neutralization and escape (as measured by PRNT titers) is only one component of the immune response to DENV. Although analysis of a longitudinal cohort study shows that these neutralization titers correlate with protection from severe secondary infection, it is unclear how PRNT titers correspond to antibody-dependent enhancement (Katzelnick et al., 2016). It is also important to note that DENV case outcomes are partially mediated by interactions with innate and T-cell immunity, the effects of which are not captured in neutralization titers (Green et al., 2014). Overall, while richer datasets and the development of more holistic assays will be required in order to fully characterize the extent of DENV antigenic diversity, it is clear that the four-serotype model is insufficient to explain DENV antigenic evolution.

### Viral clade dynamics

We use these inferred antigenic relationships to directly quantify the impact of antigenic fitness on DENV population composition. To do so, we measure serotype frequencies across Southeast Asia over time and construct a model to estimate how they will fluctuate (Methods, Eq. 6–16). This model places a fitness value on each serotype that derives from a constant intrinsic component alongside a time-dependent antigenic component. Antigenic fitness declines with population immunity, which is accumulated via the recent circulation of antigenically similar viruses. Our primary model parameterization assumes that both heterotypic and homotypic immunity wane linearly over time at the same rate, with homotypic immunity starting from a higher baseline of protection based on closer antigenic distances. We compared this to a secondary model parameterization with only heterotypic waning (see Methods), under which we observe similar model performance (Table S3).

We find that antigenic fitness is able to explain much of the observed variation in serotype growth and decline (Figure 5). Forward simulations under the optimized parameter set display damped oscillations around the serotype-specific ‘set points’ determined by intrinsic fitnesses, but intrinsic fitness alone is unable to explain serotype fluctuations (*R*^2^ = 0.04; Table S3, Figure S6). This demonstrates that although intrinsic fitness plays an important role in dictating long-term dynamics, wherein particular serotypes tend to circulate at low frequency (e.g., DENV4) and others at high frequency (e.g., DENV1 and DENV2), antigenic fitness plays out on shorter-term time scales, dictating circulation over several subsequent years.

We similarly use this model to quantify the effect of within-serotype antigenic variation on the success and decline of canonical DENV genotypes (Figure 6). As above, genotype antigenic fitness declines with population immunity. Here, we estimate population immunity based on antigenic distance from recently circulating genotypes, using distances that are either genotype-specific or based only on the serotype that each genotype belongs to. We then directly compare how strongly these coarser serotype-level versus specific genotype-level antigenic relationships impact DENV population dynamics. Overall, we find that antigenic fitness explains a moderate portion of the observed variation in genotype growth and decline. Surprisingly, however, we find that incorporating within-serotype antigenic differences does not improve our predictions (Figure 6C-D). This suggests that although genotypes are antigenically diverse, these differences do not appear to influence large-scale regional dynamics over time. This lack of signal could be explained by either (A) genotype-level frequency trajectories estimated from public data are overly noisy for this application or (B) our model of antigenic fitness based on PRNT assay data does not match reality, due to either PRNT assay data not well reflecting human immunity or due to our particular model formulation that parameterizes immunity from titer distances (Eq. 6–10). In the present analysis, we are not able to firmly resolve these disparate possibilities.

This observation is also subject to caveats imposed by the available data and model assumptions. Our estimates of antigenic fitness are informed by the antigenic distances inferred by the substitution model; thus, as above, we are unable to account for nuanced antigenic differences between sub-genotype clades of DENV due to limited titer data. We estimate DENV population composition over time based on available sequence data, pooled across all of Southeast Asia (Methods, Eq. 4). As the vast majority of cases of DENV are asymptomatic, sequenced viruses likely represent a biased sample of more severe cases from urban centers where patients are more likely to seek and access care. We also assume that Southeast Asia represents a closed viral population with homogeneous mixing. However, increasing globalization likely results in some amount of viral importation that is not accounted for in this model (Allicock et al., 2012). Finally, although Southeast Asia experiences hyperendemic DENV circulation, the majority of DENV transmissions are hyper-local (Salje et al., 2017), and viral populations across this broad region may not mix homogeneously each season. Thus, it is possible that these sub-serotype antigenic differences impact finer-scale population dynamics, but we lack the requisite data to examine this hypothesis.

## Conclusions

We find that within-serotype antigenic evolution helps explain observed patterns of cross-neutralization among dengue genotypes. We also find that population immunity is a strong determinant of the composition of the DENV population across Southeast Asia, although this is putatively driven by coarser, serotype-level antigenic differences. As richer datasets become available, future studies that similarly combine viral genomics, functional antigenic characterization, and population modeling have great potential to improve our understanding of how DENV evolves antigenically and moves through populations.

### Model sharing and extensions

We have provided all code, configuration files and datasets at github.com/blab/dengue-antigenic-dynamics, and wholeheartedly encourage other groups to adapt and extend this framework for further investigation of DENV antigenic evolution and population dynamics.

## Methods

### Data

#### Titers

Antigenic distance between pairs of viruses *i* and *j* is experimentally measured using a neutralization titer, which measures how well serum drawn after infection with virus *i* is able to neutralize virus *j in vitro* (Russell and Nisalak, 1967). Briefly, two-fold serial dilutions of serum *i* are incubated with a fixed concentration of virus *j*. Titers represent the lowest serum concentration able to neutralize 50% of virus, and are reported as the inverse dilution. We used two publicly available plaque reduction neutralization titer (PRNT50) datasets generated by Katzelnick *et al*. in (Katzelnick et al., 2015). The primary dataset was generated by infecting each of 36 non-human primates with a unique strain of DENV. NHP sera was drawn after 12 weeks and titered against the panel of DENV viruses. The secondary dataset was generated by vaccinating 31 human trial participants with a monovalent component of the NIH DENV vaccine. Sera was drawn after 6 weeks and titered against the same panel of DENV viruses. As discussed in Katzelnick *et al*., these two datasets show similar patterns of antigenic relationships between DENV viruses. In total, our dataset includes 1182 measurements across 48 virus strains: 36 of these were used to generate serum, and 47 were used as test viruses.

#### Sequences

For the titer model analysis, we used the full sequence of *E* (envelope) from the 48 strains in the titer dataset.

For the clade frequencies analysis, we downloaded all dengue genome sequences available from the Los Alamos National Lab Hemorrhagic Fever Virus Database as of March 7, 2018, that contained at least the full coding sequence of *E* (envelope) (total N=12,645) (Kuiken et al., 2011). We discarded sequences which were putative recombinants, duplicates, lab strains, or which lacked an annotated sampling location and/or sampling date. We selected all remaining virus strains that were annotated as a Southeast Asian isolate (total N = 8,644).

For both datasets, we used the annotated reference dataset from (Pyke et al., 2016) to assign sequences to canonical genotypes.

### Alignments and trees

We used MAFFT v7.305b to align nucleotide *E* gene sequences for each strain before translating the aligned sequences (no frame-shift indels were present) (Katoh and Standley, 2013). All maximum likelihood phylogenies were constructed with IQ-TREE version 1.6.8 and the GTR+I+G15 nucleotide substitution model (Nguyen et al., 2014).

### Titer Model

We compute standardized antigenic distance between virus *i* and serum *j* (denoted *D_ij_*) from measured titers *T_ij_* relative to autologous titers *T_ii_*, such that

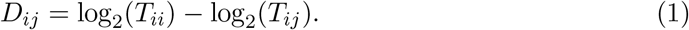

We then average normalized titers across individuals. To predict unmeasured titers, we employ the ‘substitution model’ from Neher et al. (2016) and implemented in Nextstrain (Hadfield et al., 2018), which assumes that antigenic evolution is driven by underlying genetic evolution.

In the substitution model, observed titer drops are mapped to mutations between each serum and test virus strain after correcting for overall virus avidity, *v_i_*, and serum potency, *p_j_* (‘row’ and ‘column’ effects, respectively), so that

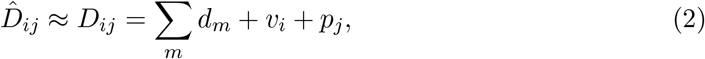

where *d_m_* is the titer drop assigned to each mutation, *m*, between serum *i* and virus *j*, and *m* iterates over mutations. We randomly withhold 10% of titer measurements as a test set. We use the remaining 90% of titer measurements as a training set to learn values for virus avidity, serum potency, and mutation effects. As in Neher et al. (2016), we formulate this as a convex optimization problem and solve for these parameter values to minimize the cost function

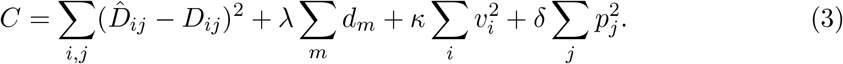

We used λ = 3.0, *κ* = 0.6, and *δ* = 1.2 to minimize test error. Respectively, these terms represent the squared training error; an L1 regularization term on mutation effects, such that most values of *d_m_* = 0; and L2 regularization terms on virus avidities and serum potencies, such that they are normally distributed. These parameter values are then used to predict the antigenic distance between all pairs of viruses, *i* and *j*. We assess performance by comparing predicted to known titer values in our test data set, and present test error (aggregated from 100-fold Monte Carlo cross-validation) throughout the manuscript.

### Viral Clade Dynamics

#### Empirical Clade Frequencies

As discussed in Neher et al. (2016) and Lee et al. (2018), we estimate empirical clade frequencies from 1970 to 2015 based on observed relative abundances of each clade in the ‘slice’ of the phylogeny corresponding to each quarterly timepoint.

Briefly, the frequency trajectory of each clade in the phylogeny is modeled according to a Brownian motion diffusion process discretized to three-month intervals. Relative to a simple Brownian motion, the expectation includes an ‘inertia’ term that adds velocity to the diffusion and the variance includes a term *x*(1 – *x*) to scale variance according to frequency following a Wright-Fisher population genetic process. This results in the diffusion process

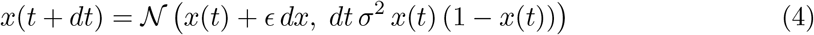

with ‘volatility’ parameter *σ*^2^ and inertia parameter *∊*. The term *dx* is the increment in the previous timestep, so that *dx* = *x*(*t*) – *x*(*t* – *dt*). We used *∊* = 0.7 and *σ* = 2.0 to maximize fit to empirical trajectory behavior.

We also include an Bernoulli observation model for clade presence / absence among sampled viruses at timestep *t*. This observation model follows

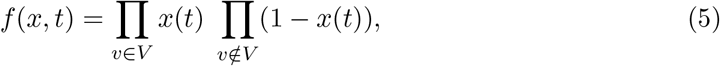

where *v* ∈ *V* represents the set of viruses that belong to the clade and *v* ∉ *V* represents the set of viruses that do not belong to the clade. Each frequency trajectory is estimated by simultaneously maximizing the likelihood of the process model and the likelihood of the observation model via adjusting frequency trajectory 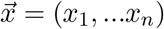.

#### Population Immunity

For antigenically diverse pathogens, antigenic novelty represents a fitness advantage (Lipsitch and O’Hagan, 2007). This means that viruses that are antigenically distinct from previously-circulating viruses are able to access more susceptible hosts, allowing the antigenically novel lineage to expand. We adapt a simple deterministic model from Łuksza and Lässig (2014) to directly quantify dengue antigenic novelty and its impact on viral fitness. We quantify population immunity to virus *i* at time *t*, *P_i_*(*t*), as a function of which clades have recently circulated in the past N years, and how antigenically similar each of these clades is to virus *i*, so that

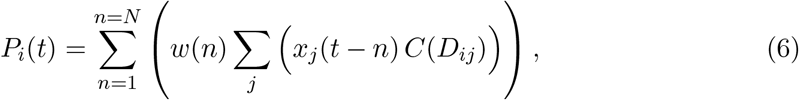

where *D_ij_* is the antigenic distance between *i* and each non-overlapping clade *j*, *n* is the number of years since exposure, and *x_j_* (*t* – *n*) is the relative frequency of *j* at year *t* – *n*. Waning immunity is modeled as a non-negative linear function of time following

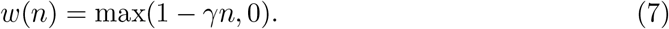

The relationship between antigenic distance and the probability of protection, *C*, is also assumed to be non-negative and linear with slope −*σ*, such that

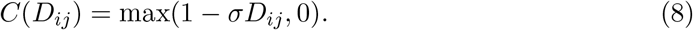

In addition to this primary analysis, we conducted a secondary analysis with a different parameterization of immunity that removes waning of homotypic immunity while allowing waning of heterotypic immunity. In this case, we assume the relationship between antigenic distance and the probability of protection, *C*, to be 50% at antigenic distance 1/*σ* and to wane based on years since infection *n* modified by *γ*_het_ following

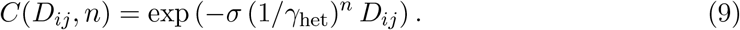

We model the effects of population immunity, *P_i_*(*t*), on viral antigenic fitness, *f_i_*(*t*), as

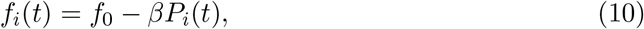

where *β* and *f*_0_ are fit parameters representing the slope of the linear relationship between immunity and fitness, and the intrinsic relative fitness of each serotype, respectively.

#### Frequency Predictions

Similar to the model implemented in Łuksza and Lässig (2014), we estimate predicted clade frequencies at time *t* + *dt* as

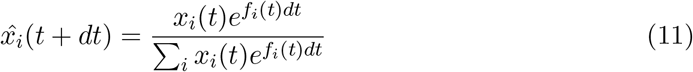

for short-term predictions (where *dt* < 1 year).

We do not attempt to predict future frequencies for clades with *x_i_*(*t*) < 0.05.

For long-term predictions, we must account for immunity accrued at each intermediate timepoint between *t* and *dt*. We divide the interval between *t* and *dt* into a total of *U* 3 month timepoints, [*t* + *u*, *t* + 2*u*,…, *t* + *U*], such that *t* + *U* = *dt*. We then compound immunity based on predicted clade frequencies at each intermediate timepoint following

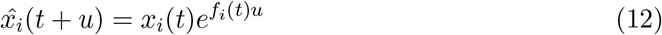

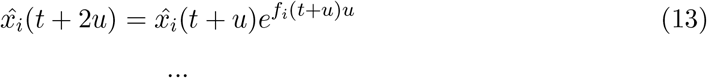

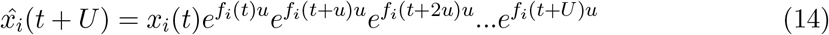

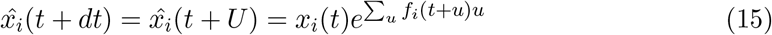

We then calculate clade growth rates, defined as the fold-change in relative clade frequency between time *t* and time *t* + *dt*

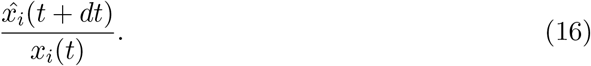

#### Null models

To quantify the impact of antigenic fitness on DENV clade success, we compare our antigenically-informed model to two null models.

Under the ‘equal fitness null’ model, all viruses have equal total fitness (antigenic and intrinsic fitness) at all timepoints

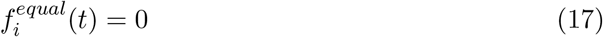

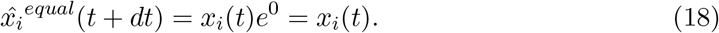

Under the ‘intrinsic fitness null’ model, all viruses have equal antigenic fitness but serotype-specific intrinsic fitness at all timepoints

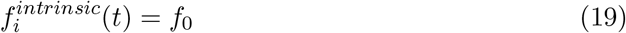

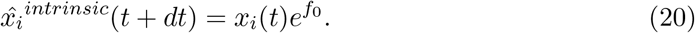

#### Model performance assessment and parameter fitting

We assess predictive power as the root mean squared error between predicted and empirical clade frequencies. To assess both the final frequency predictions and the predicted clade trajectories, this RMSE includes error for each clade, for each starting timepoint *t*, and for each intermediate predicted timepoint *t* + *u*.

Our frequency prediction model has a total of 6 free parameters. We jointly fit these parameters to minimize RMSE of serotype frequency predictions via the Nelder-Mead algorithm as implemented in SciPy v.1.0.0 (Table 1) (Gao and Han, 2012; Jones et al., 2001). We use *N* = 2 years of previous immunity that contribute to antigenic fitness and project *dt* = 2 years in the future when predicting clade frequencies.

#### Simulations

To ensure the model machinery functions correctly, we seeded a forward simulation of clade dynamics with two years of empirical frequencies and simulated predicted dynamics over the remainder of the time course (Figure S5). We then fit model parameters as described above, and obtained parameter values that well recover input values (Table 2).

**Table 2.** Parameter recovery against simulated data.

| Parameter | Input value | Optimized value |
| --- | --- | --- |
| $\beta$ | 3.25 | 3.10 |
| $\gamma$ | 0.55 | 0.56 |
| $\sigma$ | 2.35 | 2.57 |
| $f_0^{(1)}$ | 0.70 | 0.72 |
| $f_0^{(2)}$ | 0.85 | 0.78 |
| $f_0^{(3)}$ | 0.40 | 0.41 |

## Data and software availability

Sequence and titer data, as well as all code used for analyses and figure generation, is publicly available at github.com/blab/dengue-antigenic-dynamics. Our work relies upon many open source Python packages and software tools, including iPython (Pérez and Granger, 2007), Matplotlib (Hunter, 2007), Seaborn (Waskom, 2017), Pandas (McKinney et al., 2010), CVXOPT (Andersen et al., 2013), NumPy (Gao and Han, 2012; Van Der Walt et al., 2011), Biopython (Cock et al., 2009), SciPy (Jones et al., 2001), Statsmodels (Seabold and Perktold, 2010), Nextstrain (Hadfield et al., 2018), MAFFT (Katoh and Standley, 2013), and IQ-TREE (Nguyen et al., 2014). Package versions are documented in the GitHub repository.

## Acknowledgements

We would like to thank Richard Neher, John Huddleston, Andrew Rambaut, Molly OhAinle, David Shaw, Paul Edlefsen, Michal Juraska, and all members of the Bedford Lab for useful discussion and advice. SB is a Graduate Research Fellow and is supported by NSF DGE-1256082. TB is a Pew Biomedical Scholar and is supported by NIH R35 GM119774-01. LK is supported by NIH awards R01AI114703-01 and P01AI106695. Our work depends on open data sharing and many open source software tools. We gratefully acknowledge the authors and developers who make our work possible.

## Supplement

**Table S1.** Antigenically relevant mutations. Each entry represents a mutation (or colinear cluster of mutations) inferred by the titer model to have a non-zero antigenic effect size *d_m_* (shown in parentheses).

|  |  |  |
| --- | --- | --- |
| • I6V, S29G, F90Y,<br>T176P, V197I, L475M<br>(0.71) | • I139V (0.05) | • L338E (0.43) |
| • A19T (0.02) | • D154E (0.03) | • T339I (0.65) |
| • N83K (0.34) | • K160V (0.03) | • V347A (0.28) |
| • A88K (0.08) | • E161T (0.22) | • I380V (0.45) |
| • A88Q (1.39) | • A162I (0.19) | • V382A (0.04) |
| • Y90F (0.43) | • I162A (0.29) | • V382I (0.37) |
| • V91I (0.35) | • I164V (0.01) | • N385K, V454I (0.12) |
| • K93R (0.20) | • T171S (0.23) | • D390S (0.12) |
| • V114I (0.90) | • V174E (0.46) | • N390S (0.18) |
| • L122S (0.03) | • E180T (0.43) | • V454T (0.25) |
| • S122L (0.07) | • N203E (0.04) | • V461F (0.01) |
| • N124K (0.10) | • N203K (0.08) | • F461I (0.03) |
| • V129I (0.01) | • D203N (0.14) | • I462V (0.10) |
| • I129V (0.21) | • E203N (1.09) | • L462I (0.16) |
| • I129A (0.23) | • E203D (1.34) | • V462L (2.11) |
| • Y132I (0.12) | • I308V (0.18) | • T478S (0.10) |
| • Y132P, R233Q (0.34) | • G330D (0.22) | • S478M (0.43) |
|  | • I335V, N355T, P364V<br>(0.18) | • V484I (0.39) |

**Table S2.** Titer model performance comparisons. We compared performance across several different variations of the titer model. As described in Neher et al. (2016), incremental antigenic change can be assigned to either amino acid substitutions (‘Substitution’ model) or to branches in the phylogeny (‘Tree’ model). For each of these models, we can constrain the model such that antigenic change is allowed to occur only between serotypes (‘Interserotype’) or between AND within serotypes (‘Full’). For the substitution model, we constrain the interserotype model by reconstructing the amino acid sequence of the most recent common ancestor for each serotype and allowing the model to assign antigenic change only to mutations between these ancestral sequences. For the tree model, we constrain the interserotype model by allowing the model to assign antigenic change only to branches in the phylogeny that lie between serotypes. We also assess the impact of the virus avidity and serum potency terms, *v_a_* and *p_b_*. For all models and metrics, we report the mean and 95% confidence interval across 100-fold Monte Carlo cross validation with random 90%:10%, training:test splits.

| Model | Antigenic resolution | $v_a$ and $p_b$ | RMSE | Pearson $R^2$ |
| --- | --- | --- | --- | --- |
| Substitution | Full | Yes | 0.75 (0.74–0.77) | 0.78 (0.77–0.79) |
| Substitution | Full | No | 1.13 (1.11–1.16) | 0.50 (0.48–0.52) |
| Substitution | Interserotype | Yes | 0.86 (0.85–0.88) | 0.72 (0.70–0.73) |
| Substitution | Interserotype | No | 0.86 (0.84–0.87) | 0.71 (0.70–0.72) |
| Tree | Full | Yes | 0.84 (0.83–0.86) | 0.72 (0.71–0.73) |
| Tree | Full | No | 1.40 (1.38–1.42) | 0.24 (0.23–0.26) |
| Tree | Interserotype | Yes | 0.87 (0.85–0.88) | 0.70 (0.69–0.71) |
| Tree | Interserotype | No | 0.86 (0.84–0.88) | 0.72 (0.71–0.73) |

**Table S3.** Fitness model performance comparisons. Here we compare the performance of the antigenically-informed fitness models to model performance under two null formulations. In the ‘equal’ null model, all clades are assigned equal fitness (i.e., antigenic and intrinsic fitness are set to 0). In the ‘intrinsic’ null model formulation, only the serotype-specific, time-invariant intrinsic fitness values contribute to clade fitness (i.e., antigenic fitness is set to 0). For both formulations of generalized waning, all other parameters were set to the values reported in Table 1 (optimized for RMSE). Parameters for heterotypic waning were optimized separately.

| Resolution | Fitness model | Waning | RMSE | Pearson $R^2$ | Accuracy |
| --- | --- | --- | --- | --- | --- |
| Serotype | Interserotype | Generalized | 0.105 | 0.547 | 0.660 |
| Serotype | Equal fitness null | Generalized | 0.130 | 0.000 | 0.480 |
| Serotype | Intrinsic fitness null | Generalized | 0.140 | 0.042 | 0.510 |
| Genotype | Interserotype | Generalized | 0.062 | 0.286 | 0.666 |
| Genotype | Intergenotype | Generalized | 0.062 | 0.254 | 0.610 |
| Genotype | Equal fitness null | Generalized | 0.070 | 0.000 | 0.440 |
| Genotype | Intrinsic fitness null | Generalized | 0.072 | 0.032 | 0.530 |
| Serotype | Interserotype | Heterotypic | 0.109 | 0.533 | 0.666 |
| Genotype | Interserotype | Heterotypic | 0.063 | 0.291 | 0.661 |
| Genotype | Intergenotype | Heterotypic | 0.063 | 0.203 | 0.599 |

**Figure S1.**
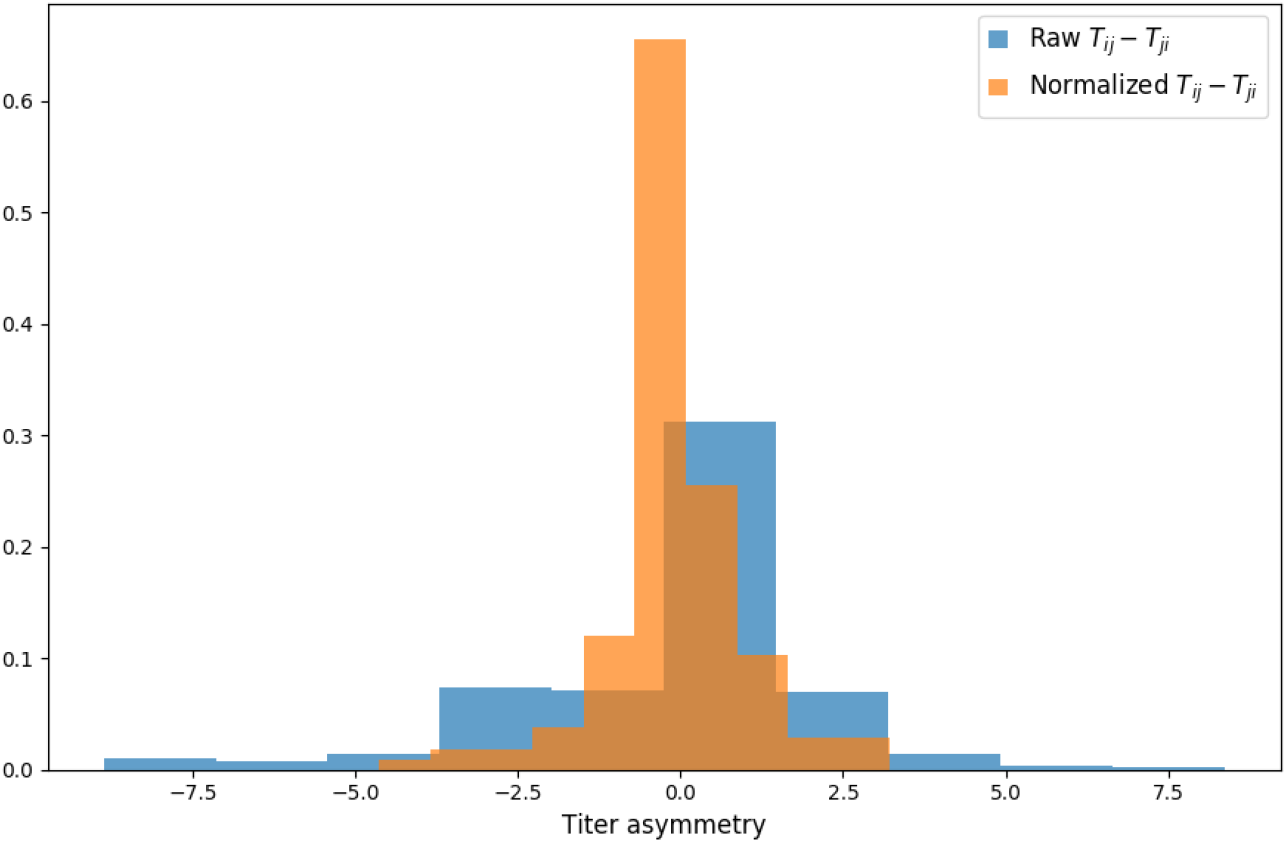
Titer value symmetry. Some viruses have greater avidity overall, and some sera are more potent overall. We normalize for these row and column effects (*v_a_* and *p_b_*, respectively) in the titer model. Once overall virus avidity and serum potency are accounted for, titers are roughly symmetric (i.e., *Dj* ≈ *Dj*).

**Figure S2.**
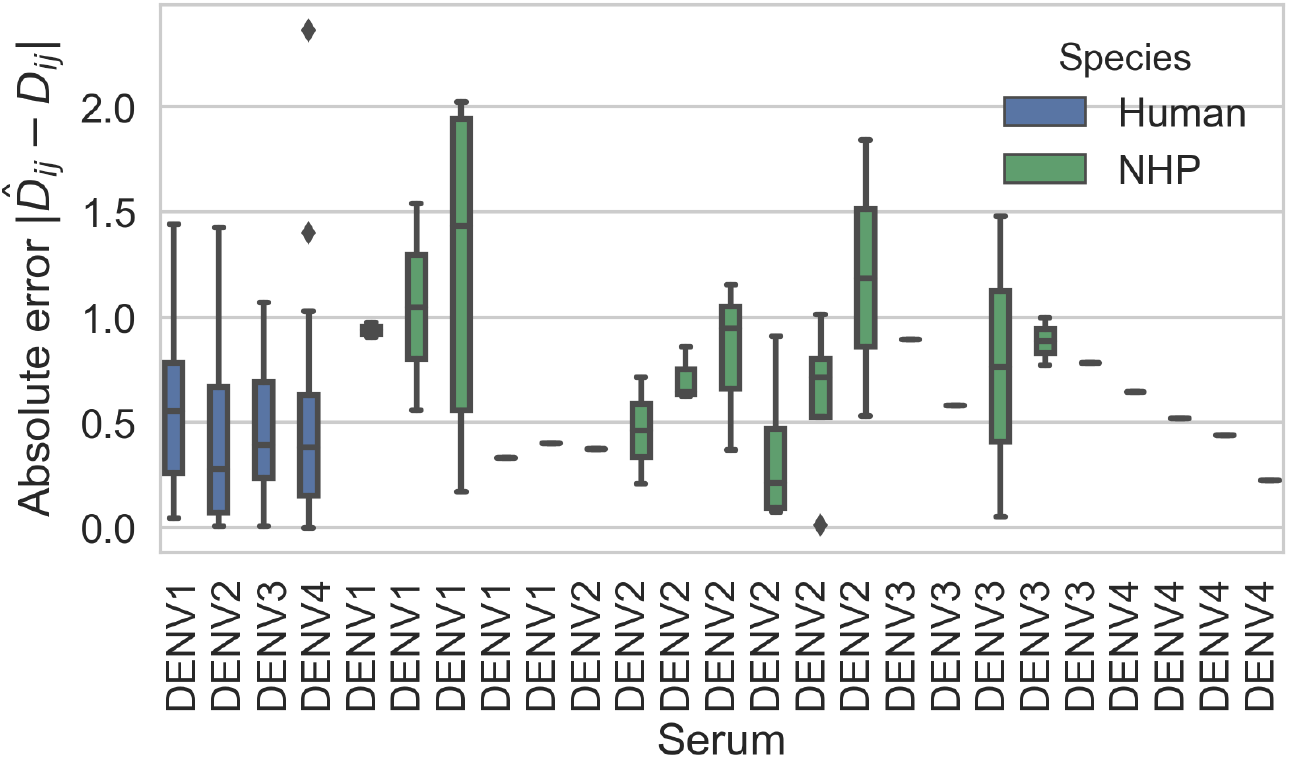
Titer prediction error by serum strain and species. Human sera was raised against four different virus strains (the monovalent vaccine components); non-human primate (NHP) sera was raised against many different virus strains. Here, we excluded NHP sera raised against the monovalent vaccine components, such that each normalized titer measurement is aggregated across individuals, but not across species. We report the out-of-sample titer prediction error for each serum strain (versus all available test viruses), aggregated across 100-fold Monte Carlo cross-validation.

**Figure S3.**
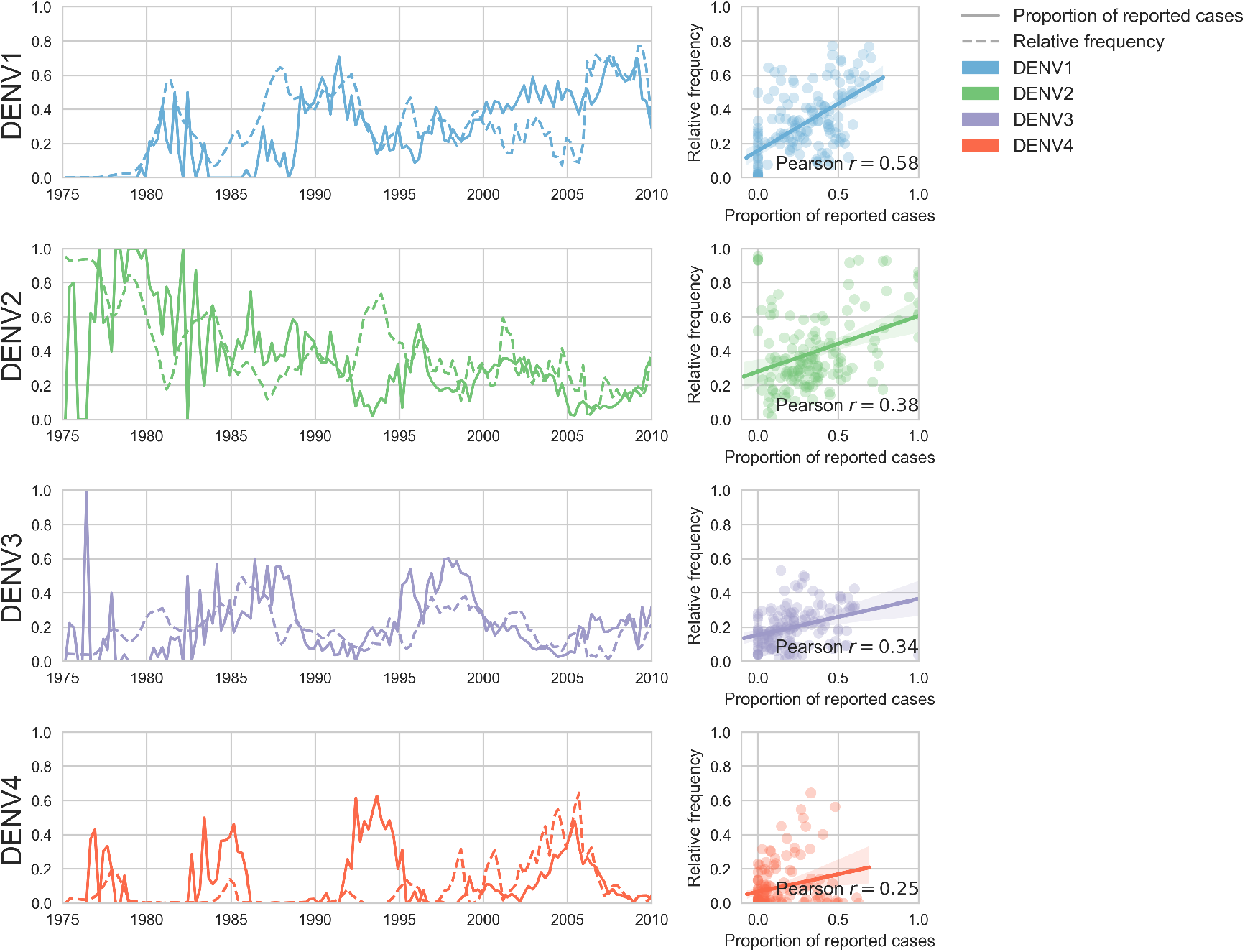
Case counts versus clade frequencies in Thailand. As described in the Methods, we estimate clade frequencies based on observed relative abundance in the ‘slice’ of the phylogeny at each quarterly timepoint. These frequency estimates are smoothed using a discretized Brownian motion diffusion process. Here, we compare estimated serotype frequencies across Thailand (all available high quality sequences) to case counts from a hospital in Bangkok between 1975–2010 (Reich et al., 2013). Biweekly case counts were aggregated into quarterly timepoints, but were not smoothed. While there are some instances where case counts and frequencies diverge (e.g., DENV4 in the early 1990s), the noisy nature of the unsmoothed case counts artificially deflates estimates of concordance.

**Figure S4.**
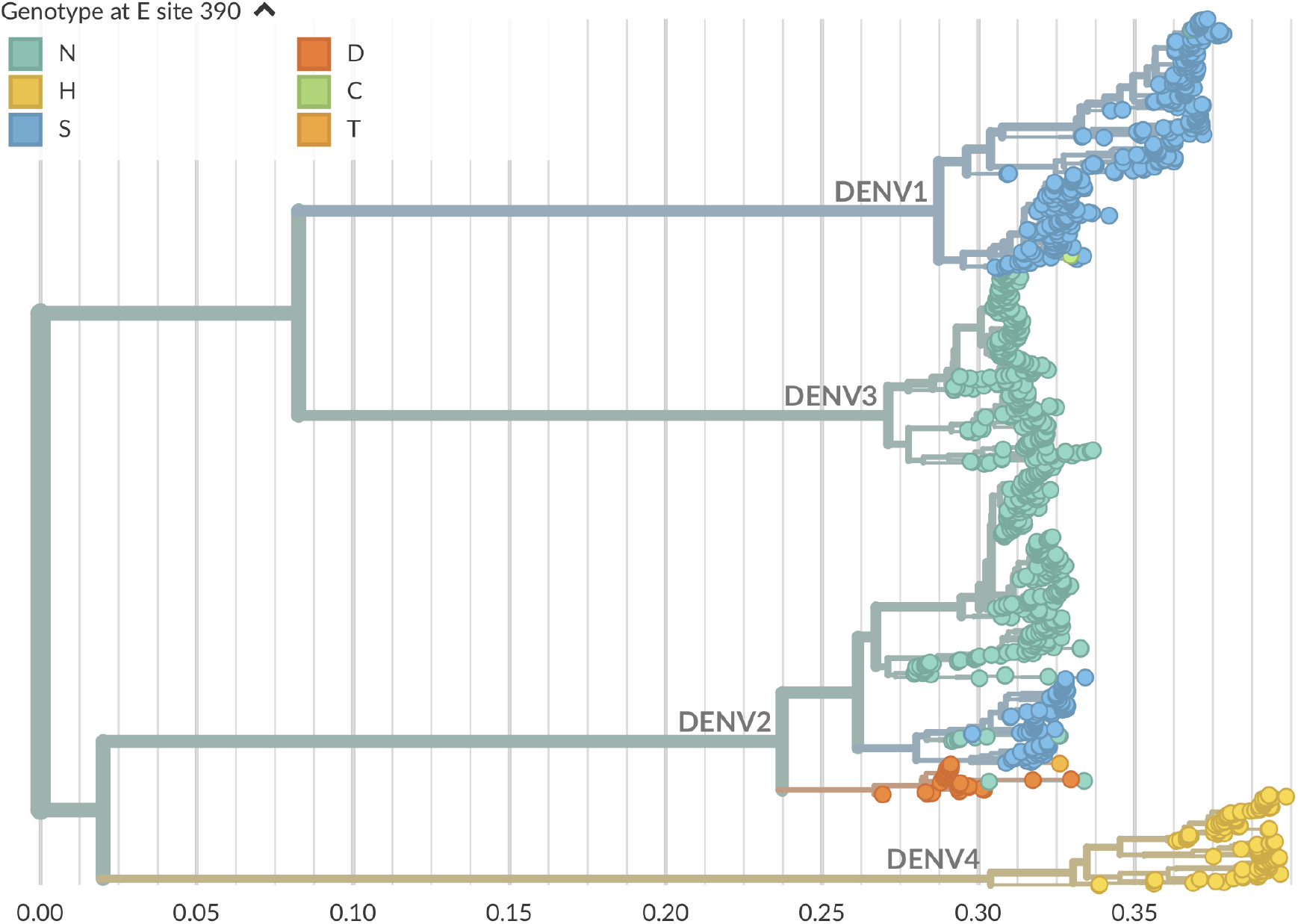
Genotype as site E 390 across dengue phylogeny. Dengue virus genotypes can be seen on Nextstrain (Hadfield et al., 2018). A live view of this figure is available at nextstrain.org/dengue/.

**Figure S5.**
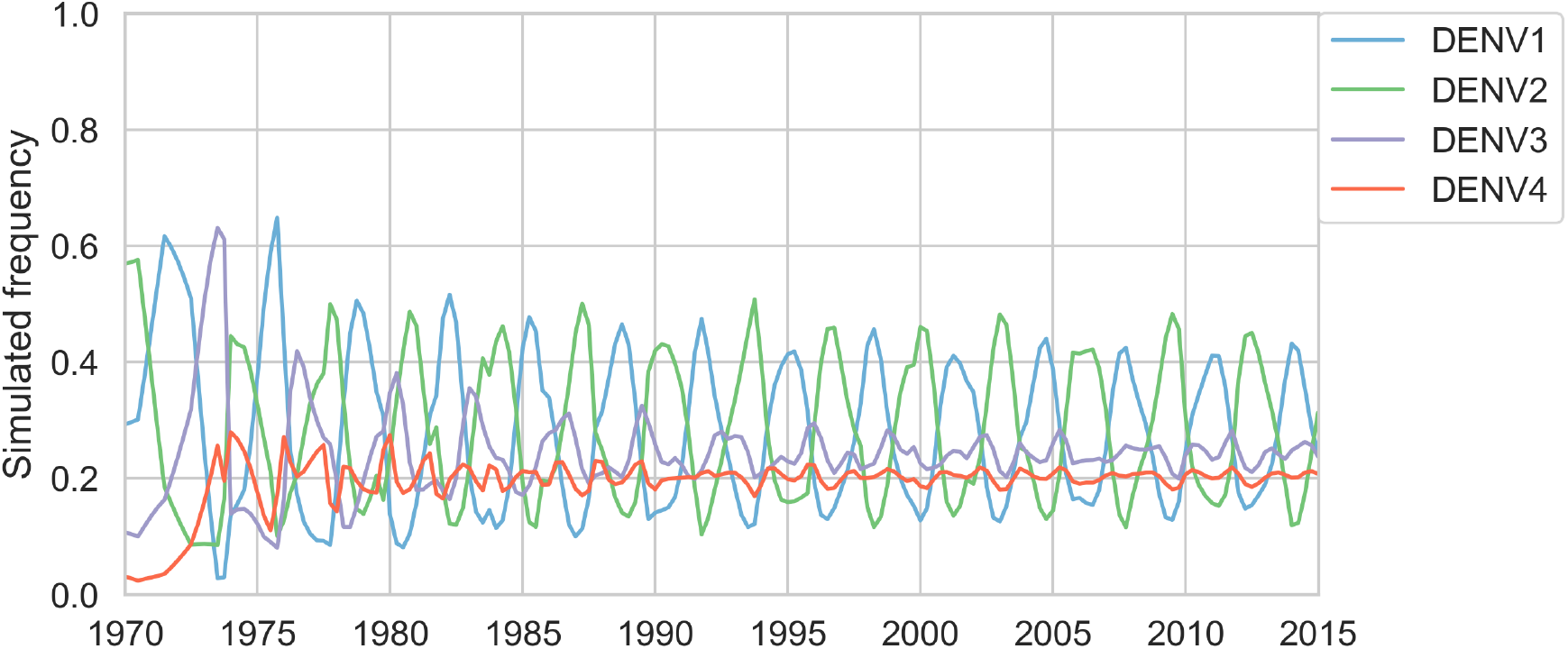
Simulated serotype frequencies. As described in the Methods, we seeded a simulation with two years of empirical frequencies and predicted forward to simulate the remainder of the timecourse. Here, we simulated under the model parameters described in Table 2

**Figure S6.**
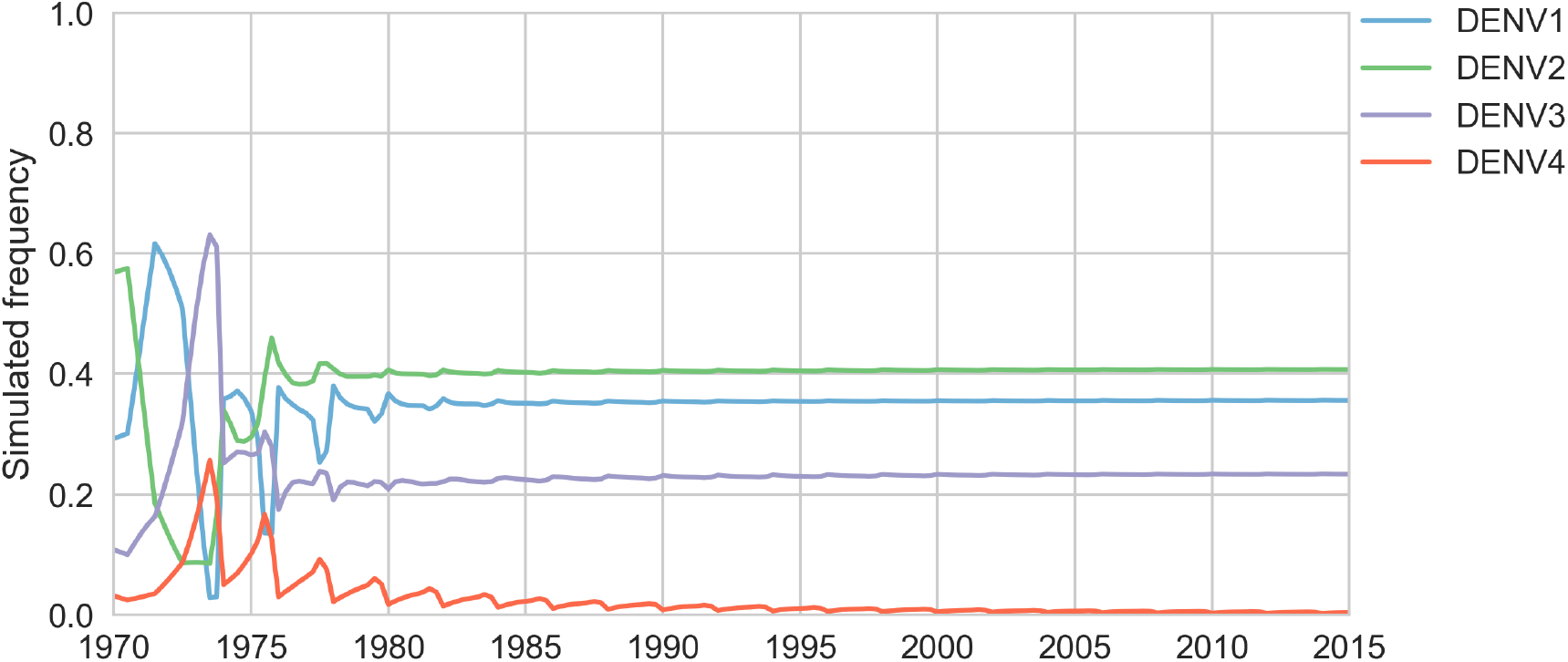
Simulated serotype frequencies (model parameters). As described in the Methods, we seeded a simulation with two years of empirical frequencies and predicted forward to simulate the remainder of the timecourse. Here, we simulated under the model parameters described in Table 1. This results in damped oscillations around the intrinsic fitness value for each serotype, but these intrinsic fitnesses alone are unable to predict observed clade dynamics (Table S3).

## References

Allicock OM, Lemey P, Tatem AJ, Pybus OG, Bennett SN, Mueller BA, Suchard MA, Foster JE, Rambaut A, Carrington CV. Phylogeography and population dynamics of dengue viruses in the Americas. Molecular biology and evolution. 2012; 29(6):1533–1543.

de Alwis R, Williams KL, Schmid MA, Lai CY, Patel B, Smith SA, Crowe JE, Wang WK, Harris E, de Silva AM. Dengue viruses are enhanced by distinct populations of serotype cross-reactive antibodies in human immune sera. PLoS pathogens. 2014; 10(10):e1004386.

Andersen M, Dahl J, Vandenberghe L. CVXOPT: A Python package for convex optimization. abel ee ucla edu/cvxopt. 2013;.

Bedford T, Cobey S, Pascual M. Strength and tempo of selection revealed in viral gene genealogies. BMC Evolutionary Biology. 2011; 11(1):220.

Bedford T, Rambaut A, Pascual M. Canalization of the evolutionary trajectory of the human influenza virus. BMC Biol. 2012; 10(1):38.

Bhatt S, Gething PW, Brady OJ, Messina JP, Farlow AW, Moyes CL, Drake JM, Brownstein JS, Hoen AG, Sankoh O, et al. The global distribution and burden of dengue. Nature. 2013; 496(7446):504.

Cock PJ, Antao T, Chang JT, Chapman BA, Cox CJ, Dalke A, Friedberg I, Hamelryck T, Kauff F, Wilczynski B, et al. Biopython: freely available Python tools for computational molecular biology and bioinformatics. Bioinformatics. 2009; 25(11):1422–1423.

Forshey BM, Reiner RC, Olkowski S, Morrison AC, Espinoza A, Long KC, Vilcarromero S, Casanova W, Wearing HJ, Halsey ES, et al. Incomplete protection against dengue virus type 2 re-infection in Peru. PLoS neglected tropical diseases. 2016; 10(2):e0004398.

Gao F, Han L. Implementing the Nelder-Mead simplex algorithm with adaptive parameters. Computational Optimization and Applications. 2012; 51(1):259–277.

Gentry M, Henchal E, McCown J, Brandt W, Dalrymple J. Identification of distinct antigenic determinants on dengue-2 virus using monoclonal antibodies. The American journal of tropical medicine and hygiene. 1982; 31(3):548–555.

Gibbons RV, Kalanarooj S, Jarman RG, Nisalak A, Vaughn DW, Endy TP, Mammen MP Jr, Srikiatkhachorn A. Analysis of repeat hospital admissions for dengue to estimate the frequency of third or fourth dengue infections resulting in admissions and dengue hemorrhagic fever, and serotype sequences. The American journal of tropical medicine and hygiene. 2007; 77(5):910–913.

Green AM, Beatty PR, Hadjilaou A, Harris E. Innate immunity to dengue virus infection and subversion of antiviral responses. Journal of molecular biology. 2014; 426(6):1148–1160.

Gupta S, Ferguson N, Anderson R. Chaos, persistence, and evolution of strain structure in antigenically diverse infectious agents. Science. 1998; 280(5365):912–915.

Hadfield J, Megill C, Bell SM, Huddleston J, Potter B, Callender C, Sagulenko P, Bedford T, Neher RA. Nextstrain: Real-time tracking of pathogen evolution. Bioinformatics. 2018; 34:4121–4123.

Halstead SB. In vivo enhancement of dengue virus infection in rhesus monkeys by passively transferred antibody. Journal of Infectious Diseases. 1979; 140(4):527–533.

Holmes EC, Twiddy SS. The origin, emergence and evolutionary genetics of dengue virus. Infection, genetics and evolution. 2003; 3(1):19–28.

Hunter JD. Matplotlib: A 2D graphics environment. Computing In Science & Engineering. 2007; 9(3):90–95. doi: 10.1109/MCSE.2007.55.

Jones E, Oliphant T, Peterson P, et al., SciPy: Open source scientific tools for Python; 2001. http://www.scipy.org/, [Online; accessed ¡today¿].

Juraska M, Magaret CA, Shao J, Carpp LN, Fiore-Gartland AJ, Benkeser D, Girerd-Chambaz Y, Langevin E, Frago C, Guy B, et al. Viral genetic diversity and protective efficacy of a tetravalent dengue vaccine in two phase 3 trials. Proceedings of the National Academy of Sciences. 2018; p. 201714250.

Katoh K, Standley DM. MAFFT multiple sequence alignment software version 7: improvements in performance and usability. Molecular biology and evolution. 2013; 30(4):772–780.

Katzelnick LC, Fonville JM, Gromowski GD, Arriaga JB, Green A, James SL, Lau L, Montoya M, Wang C, VanBlargan LA, et al. Dengue viruses cluster antigenically but not as discrete serotypes. Science. 2015; 349(6254):1338–1343.

Katzelnick LC, Gresh L, Halloran ME, Mercado JC, Kuan G, Gordon A, Balmaseda A, Harris E. Antibody-dependent enhancement of severe dengue disease in humans. Science. 2017; 358(6365):929–932.

Katzelnick LC, Montoya M, Gresh L, Balmaseda A, Harris E. Neutralizing antibody titers against dengue virus correlate with protection from symptomatic infection in a longitudinal cohort. Proceedings of the National Academy of Sciences. 2016; 113(3):728–733.

Kochel TJ, Watts DM, Halstead SB, Hayes CG, Espinoza A, Felices V, Caceda R, Bautista CT, Montoya Y, Douglas S, et al. Effect of dengue-1 antibodies on American dengue-2 viral infection and dengue haemorrhagic fever. The Lancet. 2002; 360(9329):310–312.

Kuiken C, Thurmond J, Dimitrijevic M, Yoon H. The LANL hemorrhagic fever virus database, a new platform for analyzing biothreat viruses. Nucleic acids research. 2011; 40(D1):D587–D592.

Lanciotti RS, Gubler DJ, Trent DW. Molecular evolution and phylogeny of dengue-4 viruses. Journal of General Virology. 1997; 78(9):2279–2284.

Lee JM, Huddleston J, Doud MB, Hooper KA, Wu NC, Bedford T, Bloom JD. Deep mutational scanning of hemagglutinin helps predict evolutionary fates of human H3N2 influenza variants. Proceedings of the National Academy of Sciences. 2018;.

Lipsitch M, O’Hagan JJ. Patterns of antigenic diversity and the mechanisms that maintain them. Journal of the Royal Society Interface. 2007; 4(16):787–802.

Lourenço J, Recker M. Natural, persistent oscillations in a spatial multi-strain disease system with application to dengue. PLoS computational biology. 2013; 9(10):e1003308.

Lourenço J, Tennant W, Faria NR, Walker A, Gupta S, Recker M. Challenges in dengue research: A computational perspective. Evolutionary applications. 2018; 11(4):516–533.

Łuksza and M, Lässig M. A predictive fitness model for influenza. Nature. 2014; 507(7490):57.

McKinney W, et al. Data structures for statistical computing in python. In: Proceedings of the 9th Python in Science Conference, vol. 445 Austin, TX; 2010. p. 51–56.

Mizumoto K, Ejima K, Yamamoto T, Nishiura H. On the risk of severe dengue during secondary infection: a systematic review coupled with mathematical modeling. Journal of vector borne diseases. 2014; 51(3):153.

Neher RA, Bedford T, Daniels RS, Russell CA, Shraiman BI. Prediction, dynamics, and visualization of antigenic phenotypes of seasonal influenza viruses. Proceedings of the National Academy of Sciences. 2016; 113(12):E1701–E1709.

Nguyen LT, Schmidt HA, von Haeseler A, Minh BQ. IQ-TREE: a fast and effective stochastic algorithm for estimating maximum-likelihood phylogenies. Molecular biology and evolution. 2014; 32(1):268–274.

OhAinle M, Balmaseda A, Macalalad AR, Tellez Y, Zody MC, Saborío S, Nuñez A, Lennon NJ, Birren BW, Gordon A, et al. Dynamics of dengue disease severity determined by the interplay between viral genetics and serotype-specific immunity. Science translational medicine. 2011; 3(114):114ra128–114ra128.

Olkowski S, Forshey BM, Morrison AC, Rocha C, Vilcarromero S, Halsey ES, Kochel TJ, Scott TW, Stoddard ST. Reduced risk of disease during postsecondary dengue virus infections. The Journal of infectious diseases. 2013; 208(6):1026–1033.

Pérez F, Granger BE. IPython: a system for interactive scientific computing. Computing in Science & Engineering. 2007; 9(3):21–29.

Pyke AT, Moore PR, Taylor CT, Hall-Mendelin S, Cameron JN, Hewitson GR, Pukallus DS, Huang B, Warrilow D, Van Den Hurk AF. Highly divergent dengue virus type 1 genotype sets a new distance record. Scientific reports. 2016; 6:22356.

Reich NG, Shrestha S, King AA, Rohani P, Lessler J, Kalayanarooj S, Yoon IK, Gibbons RV, Burke DS, Cummings DA. Interactions between serotypes of dengue highlight epidemiological impact of cross-immunity. Journal of The Royal Society Interface. 2013; 10(86):20130414.

Rico-Hesse R. Molecular evolution and distribution of dengue viruses type 1 and 2 in nature. Virology. 1990; 174(2):479–493.

Russell PK, Nisalak A. Dengue virus identification by the plaque reduction neutralization test. The Journal of Immunology. 1967; 99(2):291–296.

Sabin AB. Research on dengue during World War II1. The American journal of tropical medicine and hygiene. 1952; 1(1):30–50.

Salje H, Cummings DA, Rodriguez-Barraquer I, Katzelnick LC, Lessler J, Klungthong C, Thaisomboonsuk B, Nisalak A, Weg A, Ellison D, et al. Reconstruction of antibody dynamics and infection histories to evaluate dengue risk. Nature. 2018; 557(7707):719.

Salje H, Lessler J, Berry IM, Melendrez MC, Endy T, Kalayanarooj S, Atchareeya A, Chanama S, Sangkijporn S, Klungthong C, et al. Dengue diversity across spatial and temporal scales: local structure and the effect of host population size. Science. 2017; 355(6331):1302–1306.

Sangkawibha N, Rojanasuphot S, Ahandrik S, Viriyapongse S, Jatanasen S, Salitul V, Phanthumachinda B, Halstead SB. Risk factors in dengue shock syndrome: a prospective epidemiologic study in Rayong, Thailand. American journal of epidemiology. 1984; 120(5):653–669.

Seabold S, Perktold J. Statsmodels: Econometric and statistical modeling with python. In: 9th Python in Science Conference; 2010.

Stanaway JD, Shepard DS, Undurraga EA, Halasa YA, Coffeng LE, Brady OJ, Hay SI, Bedi N, Bensenor IM, Castaneda-Orjuela CA, et al. The global burden of dengue: an analysis from the Global Burden of Disease Study 2013. The Lancet infectious diseases. 2016; 16(6):712–723.

Tibshirani R. Regression shrinkage and selection via the lasso. Journal of the Royal Statistical Society: Series B (Methodological). 1996; 58(1):267–288.

Twiddy SS, Holmes EC, Rambaut A. Inferring the rate and time-scale of dengue virus evolution. Molecular biology and evolution. 2003; 20(1):122–129.

Van Der Walt S, Colbert SC, Varoquaux G. The NumPy array: a structure for efficient numerical computation. Computing in Science & Engineering. 2011; 13(2):22.

Waggoner JJ, Balmaseda A, Gresh L, Sahoo MK, Montoya M, Wang C, Abeynayake J, Kuan G, Pinsky BA, Harris E. Homotypic dengue virus reinfections in Nicaraguan children. The Journal of infectious diseases. 2016; 214(7):986–993.

Waskom M, Seaborn visualization library; 2017. http://seaborn.pydata.org/.

Wearing HJ, Rohani P. Ecological and immunological determinants of dengue epidemics. Proceedings of the National Academy of Sciences. 2006; 103(31):11802–11807.

Zhang C, Mammen MP, Chinnawirotpisan P, Klungthong C, Rodpradit P, Monkongdee P, Nimmannitya S, Kalayanarooj S, Holmes EC. Clade replacements in dengue virus serotypes 1 and 3 are associated with changing serotype prevalence. Journal of virology. 2005; 79(24):15123–15130.

